# Glioblastoma induces the recruitment and differentiation of hybrid neutrophils from skull bone marrow

**DOI:** 10.1101/2023.03.24.534105

**Authors:** By Meeki Lad, Angad S. Beniwal, Saket Jain, Poojan Shukla, Jangham Jung, Sumedh S. Shah, Garima Yagnik, Husam Babikir, Alan T. Nguyen, Sabraj Gill, Jacob S. Young, Austin Lui, Diana Salha, Aaron Diaz, Manish K. Aghi

## Abstract

Tumor-associated neutrophil (TAN) effects on glioblastoma biology remain under-characterized. We show here that ‘hybrid’ neutrophils with dendritic features – including morphological complexity, expression of antigen presentation genes, and the ability to process exogenous peptide and stimulate MHCII-dependent T cell activation – accumulate intratumorally and suppress tumor growth *in vivo*. Trajectory analysis of patient TAN scRNA-seq identifies this phenotype as a polarization state which is distinct from canonical cytotoxic TANs and differentiates intratumorally from immature precursors absent in circulation. Rather, these hybrid-inducible immature neutrophils – which we identified in patient and murine glioblastomas – arise from local skull marrow. Through labeled skull flap transplantation and targeted ablation, we characterize calvarial marrow as a potent contributor of antitumoral myeloid APCs, including hybrid TANs and dendritic cells, which elicit T cell cytotoxicity and memory. As such, agents augmenting neutrophil egress from skull marrow – such as intracalvarial AMD3100 whose survival prolonging-effect in GBM we demonstrate – present therapeutic potential.

## INTRODUCTION

Glioblastoma (GBM) is an aggressive primary brain cancer with a poor prognosis (Luo et al., 2015). Current therapies have failed in large part because they treat GBM cells in isolation and fail to account for the recent understanding that GBM is an organ with complex interplay between tumor cells and their microenvironment. GBM cells secrete several cytokines, chemokines, and growth factors, which attract the infiltration of various myeloid immune cells. These cells create a specific glioma tumor niche, which promotes tumor growth, invasiveness, and therapy resistance (Elliott et al., 2017; Gieryng et al., 2017). Three of these myeloid immune cells, tumor-associated macrophages (TAMs), tumor-associated neutrophils (TANs), and myeloid-derived suppressor cells (MDSCs) constitute the majority of nonmalignant cells in the GBM microenvironment (Badie and Schartner, 2000; Liang et al., 2014; Raychaudhuri et al., 2015).

While considerable attention has been paid to TAMs and MDSCs, little GBM research has focused on TANs even though neutrophils are the most abundant leukocytes in blood. Like macrophages, neutrophils are part of the non-specific immune system. In fact, neutrophils are the first line of defense during inflammation and infections, arriving at these sites ahead of macrophages.

Besides their classical antimicrobial functions, neutrophils are also found in many types of tumors. Early studies suggested that TANs were bystanders either being detected intravascularly as they pass through the tumor or passively accumulating in the tumor without actual function because it was hard to imagine that neutrophils, being short-lived cells, could have an effect on chronic and progressive diseases such as cancer. More recent studies have cast doubts on this premise by suggesting that TANs influence tumor cells and cells in the microenvironment, although their roles vary considerably and can be pro-tumoral (Liang et al., 2014) or anti-tumoral (Singhal et al., 2016). In GBM, the finding that an elevated peripheral neutrophil-to-lymphocyte ratio (NLR) is associated with poor survival in GBM patients (Bambury et al., 2013) has led some to favor a pro-tumor role for TANs, but these correlative studies of circulating peripheral blood neutrophils (PBNs) have yet to be supplemented with mechanistic studies of TANs and have not determined the interaction between neutrophils and lymphocytes, the two components of the NLR, in GBM. The tumor-specific cues that dictate TAN functionality and the nature of TANs in GBM, which has already been described as a T cell-depleted (Chongsathidkiet et al., 2018) and macrophage-rich (Badie and Schartner, 2001; Muller et al., 2017) tumor, thus remain unclear.

To address this knowledge gap, we isolated PBNs and TANs from GBM patients, compared their transcriptomic profiles, analyzed their interactions with tumor cells and T-cells in culture, and determined their effects *in vivo* in the presence or absence of T-cells. We then used trajectory analysis from single cell RNA-sequencing of patient PBNs and TANs and a novel calvarial transplantation technique to define a distinct PBN-independent lineage of patient TANs from immature neutrophil precursors in the skull bone marrow.

## RESULTS

### TANs are durable residents of the perivascular niche of GBM

Flow cytometric characterization of newly diagnosed GBM patient samples revealed that CD45^+^CD16^+^CD66b^+^ TANs constituted 0.39% [range: 0.11-0.69%] of total tumor cellularity and 4.04% [range 2.52-7.78%] of tumor-infiltrating leukocytes (n=6; **Fig. 1A; Supp. Table 1**). To determine if these neutrophils were more than passerby intravascular cells, we performed IHC to co-localize MPO^+^ TANs and blood vessels in patient GBMs. We found that MPO^+^ TANs indeed resided in the GBM parenchyma, approximating blood vessels within the perivascular niche but not occupying the intravascular compartment (**Fig. 1B**).

**Figure 1.**
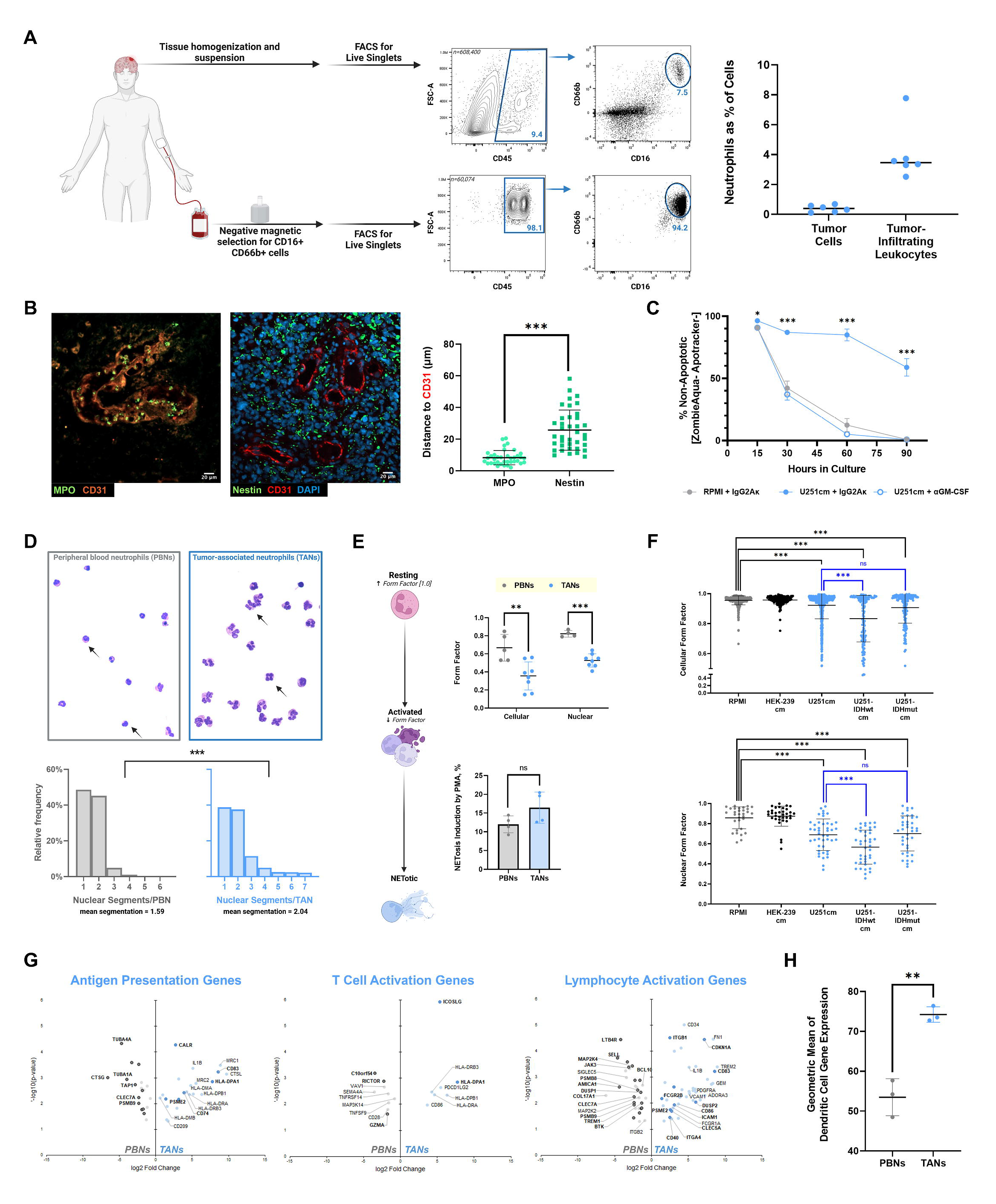
TANs are robust, morphologically activated residents of the perivascular niche in GBM. **(A)** Schematic depicting patient TAN and PBN isolation, with TAN infiltration quantified as a percentage of both total tumor cells and tumor-infiltrating leukocytes (n=6). **(B)** Immunohistochemical comparison of endothelial (CD31^+^) proximity to canonically perivascular glioma stem cells (Nestin^+^) versus neutrophils (MPO^+^) in human GBM (n=37-38 cells from 3 GBMs) **(C)** The effect of U251cm on longitudinal neutrophil survival compared to control media (P_15h_=0.03, P_30-90h_<0.001), and its neutralization by GM-CSF blockade (n=6/group). **(D)** Nuclear segmentation of Wright-Giemsa stained cytospins of patient TANs and PBNs. **(E)** *Upper panel:* Comparison of membrane circularity (form factor) at the cellular and nuclear levels between DAPI-stained TANs and PBNs (P_cellular_=0.001, P_nuclear_<0.001; n=4-8/group). *Lower panel:* NETosis induction by LPS in cultured TANs and PBNs (P=0.125; n=4/group) **(F)** Effects of U251cm, either basal or derived from IDH-wt or IDH-mut overexpressing variants, on morphological activation (reflected in lower cellular and nuclear form factor) of PBNs compared to control (RPMI) or non-GBM- conditioned media (HEK239cm) conditions. N=150-743 cells/group. **(G-H)** Nanostring multiplex assay of myeloid gene expression in TANs and matched PBNs (n=3 pairs). **G**: Volcano plots depicting significantly (p<0.05) differentially expressed genes related to antigen presentation and T cell/lymphocyte activation. **H:** Geometric mean of normalized dendritic cell-related gene expression per sample. *Data are represented as means ± SD. *p<0.05, **p<0.01, ***p<0.001*

Our analysis also indicated that IDH-wild type GBMs may be preferentially chemotactic towards these neutrophils compared to IDH-mutant GBMs, as TAN infiltration was greater in the former (n=9/group; P<0.05; **Supp Fig. 1A**). This finding was recapitulated in an analysis of myeloperoxidase (MPO) expression using The Cancer Genome Atlas (TCGA) dataset (P<0.001; **Supp Fig. 1B**). To interrogate differences in chemotaxis, we compared healthy PBN migration towards tumor-conditioned media from U251 glioblastoma cells engineered to overexpress either wild-type or mutant IDH, and confirmed that the former induced greater mobilization (P<0.001; **Supp. Fig. 1C**).

Further, we found that PBN longevity was enhanced upon exposure to the GBM secretome, dispelling the notion that TANs are short-lived effectors and suggesting that, despite their general scantness, these cells may have relevant pathophysiologic properties given their endurance. Indeed, healthy control PBNs exhibited robust survival as late as 90 hours (58.83% [95% CI: 53.2-64.5%]) when grown in media conditioned by the U251 glioblastoma cell line (U251cm), as compared to almost no survival (0.9% [0.8-1.0%]) in control conditions (P=0.03 15 hrs, <0.001 30-90 hrs; **Fig. 1C; Supp. Fig. 1D**). This effect of GBM cells promoting PBN survival was completely abrogated by the administration of a GM-CSF blocking antibody (P=0.03 15 hrs, <0.001 30-90 hrs; **Fig. 1C**), suggesting that this growth factor is crucial to extended neutrophil longevity in the tumor microenvironment.

### Compared to PBNs, TANs are morphologically activated in a manner promoted by the GBM secretome

We then employed Wright-Giemsa staining to both validate the purity of our neutrophil FACS sorting algorithm and to evaluate gross morphologic differences between TANs and patient-matched PBNs. TANs had visibly greater nuclear segmentation than PBNs, consistent with neutrophil activation: 8.2% of TANs were hyper-segmented (5+ nuclear segments), compared to 0.2% of PBNs (P<0.001; **Fig. 1D**). Form factor analysis recapitulated this observation at the nuclear (P=0.0012; **Fig.1E**) and cytoplasmic (P<0.001; **Fig. 1E**) membrane levels, demonstrating that TANs had relatively deformed, non-spherical membranes consistent with nuclear segmentation and cytoplasmic flattening, respectively. Of note, these morphological changes in TANs did not impact their capacity for NETosis, the primary effector mechanism by which neutrophils expel their DNA and intracellular contents in a web-like neutrophil extracellular trap (NET). TANs and PBNs, regardless of starting differences in shape, exhibited similar NETotic capacity when stimulated by PMA (P=0.1; **Fig. 1E**). Finally, we observed that U251cm promoted this morphologic change in healthy volunteer circulating PBNs (P<0.001; **Fig. 1F**), confirming the role of the GBM secretome in mediating this phenomenon. Furthermore, IDH-wt-overexpressing U251cm caused greater overall membrane deformation (P<0.001; **Fig. 1F**) than IDH- mut-overexpressing U251cm.

### GBM TANs exhibit a non-canonical APC transcriptional signature

We then transcriptomically compared TANs to PBNs from the same GBM patients (n=3 pairs) using the NanoString nCounter platform and a multiplex panel of over 500 immunology genes. Compared to PBNs, TANs upregulated genes associated with antigen presentation, including costimulatory ligands (CD83, CD86, CD40, ICOSLG), MHCII subunits (HLA-DRB3/A, DPA1/B1), and antigen processing proteasomes and chaperones (CD74, CALR, PSME2, HLA-DMA/B) (**Fig. 1G**, **Supp. Fig. 1E, Supp. Table 2**). Many of these genes were canonical dendritic cell-related genes, consistent with previously reported neutrophil-dendritic cell “hybrids” observed in autoimmune disease (Ashtekar and Saha, 2003) and lung cancer (Singhal et al., 2016). Indeed, geometric mean gene expression of dendritic cell-related genes was higher in TANs than PBNs (P=0.002, **Fig. 1H**). Additionally, pathway analysis revealed that TANs exhibited increased expression of genes in the PI3-Akt and MAPK signaling pathways compared to PBNs (**Supp Fig. 1F**).

### The GBM secretome does not fully convert PBNs to an APC transcriptional signature

Culturing healthy volunteer PBNs in U251cm caused similar upregulation of many, though not all, of the same antigen presentation-related genes (**Supp. Fig. 1G-H, Supp. Table 3**). While part of this discontinuity was attributable to allelic MHCII variance, the inability for the GBM secretome to induce expression of the potent costimulatory ligand CD86, HLA-DM chaperones, and PSME2 suggested that healthy PBNs may be unable to fully adopt an APC-TAN phenotype. This was further supported by the fact that aggregate dendritic cell gene expression did not significantly increase with healthy PBN exposure to the GBM secretome (**Supp. Fig. 1I**).

### GBM TANs activate T-cells in an MHC-II dependent manner

We then determined if there were functional consequences of the expression of these antigen presentation genes. Flow cytometric analysis of newly diagnosed GBMs confirmed that 11.4-35.7% of TANs expressed MHC-II (HLA-DR/DP/DQ), while patient-matched PBNs did not (n=8; P<0.001; **Fig. 2A**). MHC-II^+^ TANs also exhibited greater side scatter on flow cytometry than MHC-II^-^ TANs from the same patients, a marker of the internal complexity (i.e. granularity) of a cell (n=8; P=0.004; **Fig. 2B**). Using DQ Ovalbumin, a fluorogenic substrate for proteases, we confirmed that these MHC-II^+^ TANs possessed the internal machinery necessary to process exogenous peptide antigen, unlike MHC-II^-^ TANs or PBNs (n=3/group; p<0.001; **Fig. 2C**).

**Figure 2.**
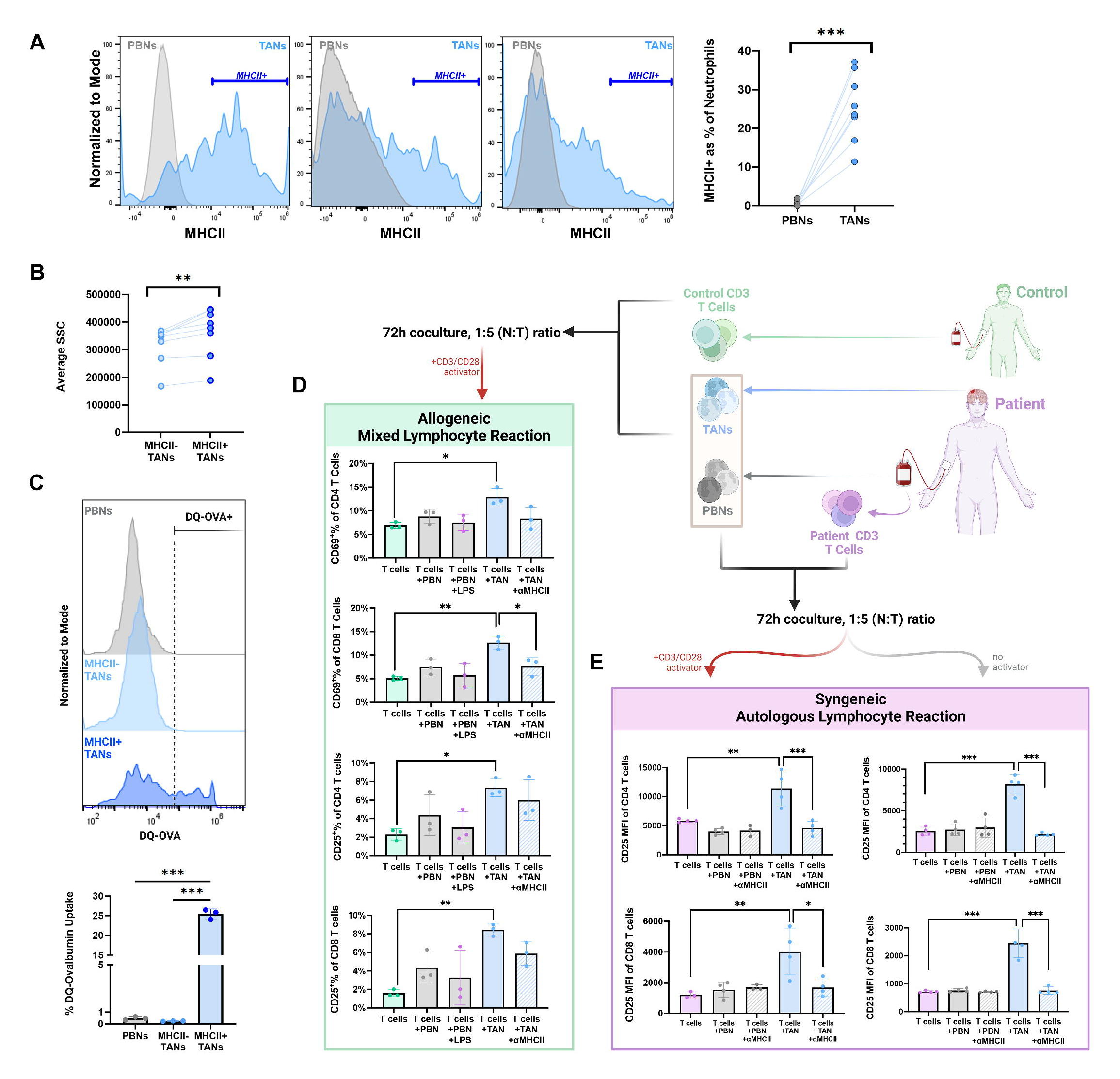
TANs activate T-cells in an MHC-II dependent manner. **(A)** Flow cytometric characterization of surface MHCII expression in GBM patient TANs and PBNs (n=8), with 3 representative patient histograms depicted. **(B)** Comparison of side scatter (SSC) between MHCII^+^ and MHCII^-^ TANs in 8 patient GBMs (P=0.004). **(C)** Representative histograms comparing exogenous antigen (DQ-OVA) uptake and processing by PBNs, MHCII^-^ TANs, and MHCII^+^ TANs from a GBM patient, with aggregate results from technical replicates depicted below (n=3/group). **(D)** Early (CD69%) and late (CD25%) activation of healthy volunteer CD3^+^ T cells cultured with patient neutrophils for 72h in the presence of 10 µL/mL CD3/CD28 activator, compared to T cells incubated in isolation. P_CD69%_ of CD4^+^=0.011, CD8^+^=0.002; P_CD25%_ of CD4^+^=0.026, CD8^+^=0.003 (n=3/group). To determine the dependence of T cell activation on stimulation by TANs, TANs were treated with 10 µg/µL αMHCII (P_CD69%_ of CD8 = 0.033). **(E)** Late activation (CD25 MFI) of patient CD3^+^ T cells cultured with autologous neutrophils for 72h, with or without 10 µL/mL CD3/CD28 activator, compared to T cells incubated in isolation. P_CD3/28-_ for TAN cocultures<0.001 CD4^+^/CD8^+^; P_CD3/28+_=0.001 CD4^+^, 0.004 CD8^+^. Antibody-mediated quenching of surface MHCII on neutrophils prior to coculture abrogated all TAN stimulatory effects (P_CD3/28+_=0.010 for CD8^+^, P<0.001 for remaining conditions). N=3/group. *Data are represented as means ± SD. *p<0.05, **p<0.01, ***p<0.001*

We then performed mixed allogenic and autologous syngeneic lymphocyte reactions to determine the effects of TANs on T-cells and the role of MHC-II in these interactions. In the mixed lymphocyte reactions (MLRs), where healthy volunteer CD3^+^ T cells were cocultured with patient TANs vs PBNs for 72 hours in the presence of a CD3/28 T cell activating cocktail, we observed that TANs enhanced early and late activation (CD69 and CD25 expression, respectively) of CD4^+^ and CD8^+^ T cells (p_CD4[early]_=0.01; p_CD4[late]_=0.03; p_CD8[early]_=0.002; p_CD8[late]_=0.003; **Fig. 2D**). Conversely, PBNs, even when primed with endotoxin, did not yield T cell activation, suggesting that the TANs are phenotypically distinct from resting and conventionally activated PBNs. In the autologous lymphocyte reactions, coculture of patient TANs with matched circulating CD3^+^ T cells from the same patients similarly yielded enhanced late activation, both with and without the addition of CD3/28 (with activator: p_CD4[late]_=0.002, p_CD8[late]_=0.004; without activator: p_CD4[late]_<0.001, p_CD8[late]_<0.001; **Fig. 2E**), suggesting that TANs express costimulatory ligands and are thus capable of independently providing the dual signals necessary for T cell activation. Administration of anti-MHCII antibody to cultures abrogated the stimulatory effect of TANs in the autologous reactions (P≤0.01; **Fig. 2E**), confirming the dependence of this phenomenon on peptide cross presentation.

### TANs retain their immunostimulatory antigen-presenting features in vivo and suppress tumor growth

We then investigated murine GBM models to discern if these models developed MHCII^+^ TANs and if these hybrid TANs accumulated with sufficient magnitude to elicit effects on tumor burden *in vivo* consistent with their immunostimulatory properties in culture. First, we confirmed in two syngeneic murine GBM models (GL261 tumors in C57BL/6J mice and BGL1 tumors in Balb/cJ mice) that overall neutrophil infiltration extent was comparable to that in human tumors (**Supp. Fig. 2A**), and that only intratumoral CD11b^+^Ly6G^+^ neutrophils appreciably expressed MHC-II (HLA-IA/E; P=0.002 BGL1 in Balb/cJ; P=0.02 GL261 in C57/BL6; **Supp. Fig 2B**). Again, analogous to our patient findings, these MHCII^+^ TANs were morphologically distinct, exhibiting higher forward and side scatter than their conventional MHC-II^-^ TANs, reflecting their greater size (P<0.001 BGL1 in Balb/cJ; P=0.004 GL261 in C57/BL6; **Supp. Fig. 2C**) and internal complexity (P=0.002 BGL1 in Balb/cJ; P<0.001 GL261 in C57/BL6; **Supp. Fig. 2C**), respectively.

We then systemically depleted neutrophils via αLy6G (clone 1A8) in BGL1 GBM-bearing Balb/cJ mice, electing this strain given its greater long-term responsiveness to depletion (Boivin et al., 2020) compared to C57BL/6J mice, in which resistance develops by 2 weeks (Magod et al., 2021). Treatment was initiated with a single intraperitoneal loading dose prior to tumor implantation and subsequent maintenance injections for 21 days. Depletion, as monitored systemically and intratumorally, was initially robust (1 week efficacy: 90.1% systemically, 95.0% intratumorally) and remained meaningfully sustained over the entire experimental course (3-week efficacy: 67.0% systemically, 49.0% intratumorally; **Supp. Fig. 3A**).

Consistent with an antitumoral TAN phenotype, tumor growth was accelerated in αLy6G-treated mice, with neutrophil-deficient tumors measuring 2.4-fold larger by bioluminescent imaging (BLI) at endpoint compared to isotype-treated controls (p=0.038; **Fig. 3A**). This increase in tumor burden was concomitant with a shift in intratumoral T cell distribution from CD8^+^ CTL-enriched (P=0.015) to double-negative T (DNT) cell-enriched (P=0.008), corroborating prior reports of neutrophil-mediated suppression of TCRγδ T cell proliferation (**Fig. 3B**) (Kalyan and Kabelitz, 2014; Mensurado et al., 2018; Oberg et al., 2019; Sabbione et al., 2014). In fact, CD8^+^ proportion correlated linearly and inversely (R^2^=0.5917, p=0.010) with tumor burden (**Fig. 3B**), suggesting that neutrophil-mediated recruitment and activation of CTLs attenuated tumor growth *in vivo*. This phenomenon was further supported by transcriptional disparities between treated and control tumors, as evaluated by a Nanostring nCounter tumor-signaling panel; neutrophil-deficient mice had lower cell type scores for cytotoxic cells in general (P<0.01), T-cells (P<0.05), and NK cells (P<0.05) (**Fig. 3C**). Moreover, CD8a expression not only correlated robustly with gross CD8^+^ T cell infiltration (R^2^=0.9519, p<0.001; **Supp. Fig. 3B**), but also with the expression of cytotoxicity-related genes (**Fig. 3D**), suggesting that these accumulated CTLs were activated effectors. These genes included the classical toxins perforin and granzyme A (R^2^ =0.5428, p=0.003; R^2^ =0.5428, p=0.015); KLRK1, which encodes activating receptor NKG2D (R^2^_KLRK1_=0.7569, p=0.001); and degranulation mediator NKG7 (R^2^=0.8991, p<0.001) (**Fig. 3D**) (Ng et al., 2020). In fact, Gene Ontology pathways spanning myriad other T cell-related functions – TCR signaling (GO 0050852, p<0.001), differentiation (GO 0045582, p<0.001), activation (GO 0046635, p=0.007), and proliferation (GO 0042102, p<0.001) - were also upregulated in CD8a^hi^ tumors compared to CD8a^lo^ ones (**Supp. Fig. 3C**).

**Figure 3.**
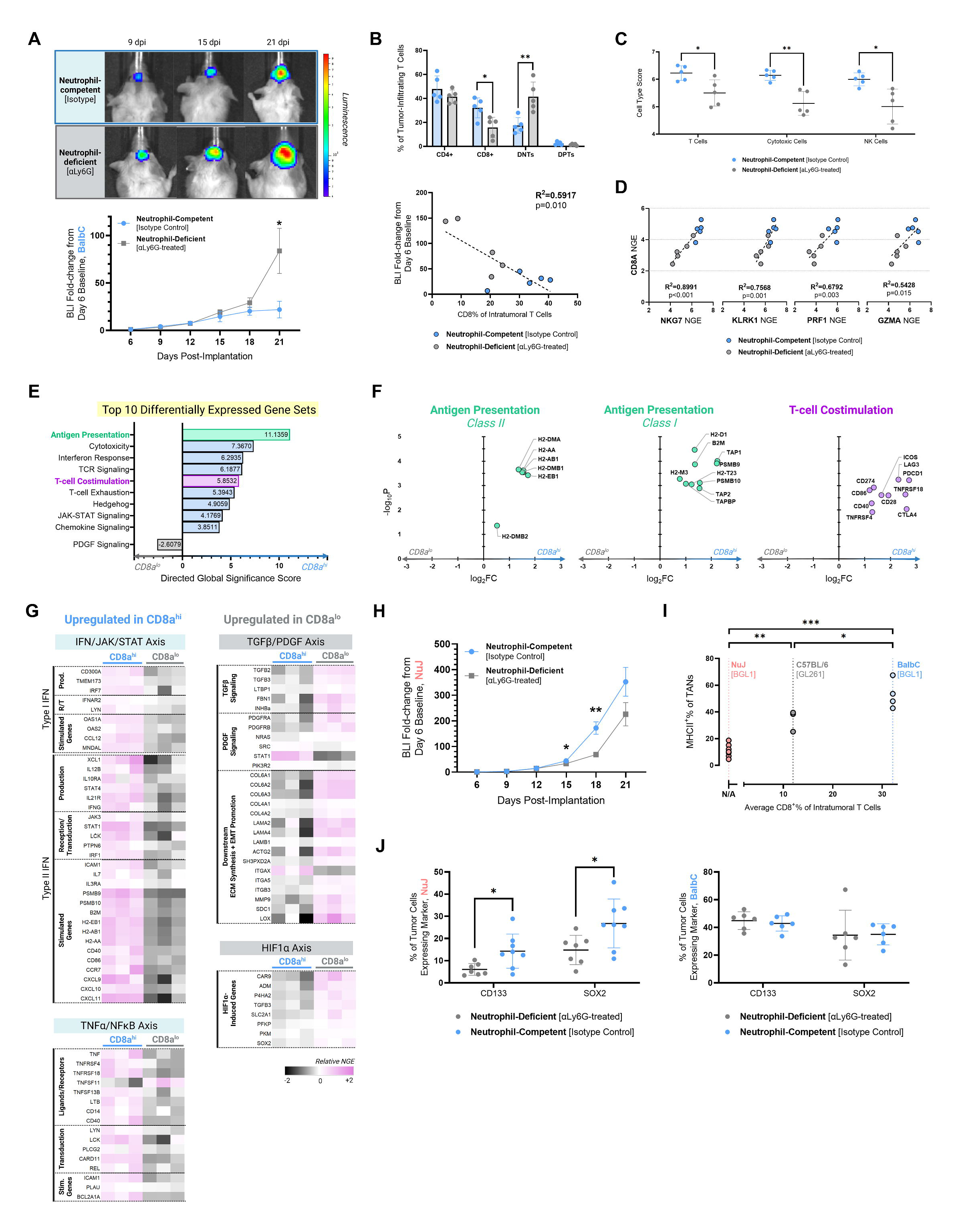
Neutrophil depletion exerts T-cell dependent effects on GBM in vivo. **(A-G)** BGL1 GBM-bearing immunocompetent BalbC/J mice were treated with αLy6G (1A8; n=9) or matched isotype (2A3; n=8) every 3 days, initiated 24h prior to tumor implantation. **A:** Tumor growth as proxied by bioluminescent imaging (BLI)^i^ (P_POD21_=0.039). Images from 1 representative mouse per cohort shown above. **B:** Flow cytometric characterization of tumor-infiltrating T cells (*above*; P_CD8_=0.015; P_DNT_=0.008), and the correlation of one subset (CD8^+^) with tumor growth as measured by total BLI fold-change. **C:** Cell type scores as derived from a Nanostring multiplex tumor signaling gene expression panel run on POD21 tumors (n=5/group). P_T_ =0.025, P_Cytotoxic_=0.003, P_NK_=0.022. **D:** Correlation of normalized gene expression (NGE) of cytotoxicity effector function genes with NGE of CD8a. **E-G:** Nanostring transcriptomic comparison of the top 3 most-and least-CD8a enriched POD21 tumors. (E) Top differentially expressed gene sets. (F) Significantly (p<0.05) differentially expressed genes related to antigen presentation and T-cell co-stimulation. (G) Heatmap of select signaling transduction cascade genes. **(H)** Tumor growth (BLI^i^) in BGL1 GBM-bearing immunodeficient Nu/J mice treated with αLy6G or matched isotype (n=9 and 10, respectively). P_POD15_=0.011, P_POD18_=0.002. **(I)** Prevalence of MHCII^+^ TANs across different murine GBM models, stratified by the model mean CD8^+^% of tumoral T cells. P_vs(Nu/J)_=0.002 for C57BL/6J, <0.001 for BalbC/J. P_vs(C57BL/6J)_=0.018 for BalbC/J. **(J)** Flow cytometric characterization of stem marker (CD133, SOX2) expression in αLy6G- vs isotype-treated mice for both models: Nu/J (*left*; P_CD133_=0.020; P_SOX2_=0.024) and BalbC/J (*right*). *Data are represented as means ± SD, except for BLI, where error bars indicate SEM*. *\*p<0.05, **p<0.01, ***p<0.001* *^i^BLI values normalized to initial POD6 recording, per mouse*

To better discern the inflammatory mediators linking TAN infiltration with CD8^+^ T cell enrichment, we compared the transcriptional profiles of the three tumors most and least enriched in CD8a expression, which came from control and αLy6G-treated mice, respectively (**Supp Table 4**). Gene set analysis demonstrated that cumulatively, CD8a^hi^ and CD8a^lo^ tumors were most stratified in their expression of antigen presentation elements (**Fig. 3E**). Beyond illustrating a connection between the hybrid TAN phenotype and CTL enrichment, the robust upregulation of these genes in CD8a^hi^ GBMs suggested that TANs – though not canonical APCs – contributed meaningfully to thi biological process in the GBM microenvironment. In fact, mean expression of 5 of 6 differentially expressed MHC class II subunits (Eb1, DMb1, Ab1, Aa, DMa, and DMb2) was over 2-fold higher in CD8a^hi^ (neutrophil-competent) mice than their CD8a^lo^ (neutrophil-deficient) counterparts (**Fig. 3F**), and similar magnitudes of upregulation were observed for MHC class I- and costimulation-related genes (**Fig. 3F**).

### IFN- and TNFα-dependent signaling are implicated in TAN-mediated CD8+ T-cell enrichment, whereas EMT- and stem-promoting PDGF/TGFβ pathways predominate in TAN-deficient GBMs

When assessing putative signaling networks involved in this TAN-CTL axis, we observed that CD8a^hi^ tumors upregulated the inflammatory IFN-JAK-STAT and TNFα- NFκB pathways (**Fig. 3G**). Upstream mediators of both type I and II interferon production were preferentially expressed in CD8a^hi^ tissues, including IL12B (log_2_FC=1.5158, p=0.027) and IL21R (log_2_FC=1.7901, p=0.001), which induce the differentiation of IFNγ-producing Th1 cells (Jorgovanovic et al., 2020; Wan et al., 2015). In turn, downstream expression of the corresponding interferon-stimulated genes (ISGs) was similarly enriched, namely the T cell chemokines CXCL9/10/11, adhesion receptor ICAM1, and elements of class I and II presentation (e.g., costimulatory ligands CCR7/CD86/CD40, PSMB proteasomes, and MHCI/II subunits). Critically, the implication that IFNγ mediated the induction of antigen-presenting features in TANs corroborated prior reports that this cytokine – alongside other circulating factors like GM-CSF, IL-3, and TNFα – is integral to the hybrid phenotype transition (Takashima and Yao, 2015). Indeed, signaling related to two of these other factors – IL3RA (log_2_FC=0.2653, p=0.049) and TNF (log_2_FC=1.1243, p=0.010) – was also upregulated in CD8a^hi^ tumors.

By contrast, HIF1α and TGFβ/PDGF axes predominated in CD8a^lo^ tumors, which converged in their transcriptional promotion of angiogenesis and GBM stem cells (GSCs). HIF1α-induced adrenomedullin (log_2_FC=-2.2799, p=0.015) and PDGFRA/B signaling (log_2_FC=-1.9310, p=0.011; log_2_FC=-1.8590, p=0.004) increase VEGF synthesis, thus enhancing vascular proliferation (Cheng et al., 2017). Meanwhile, HIF1α-induced SOX2 (log_2_FC=-1.2375, p=0.022) and TGFβ superfamily cytokine activin A (encoded by INHBa; log_2_FC=-0.8391, p=0.029) induce GBM stemness and self-renewal (Lopez-Bertoni et al., 2022; Pauklin and Vallier, 2015). This stemness phenotype likely had further contribution from the reduced IFNγ expressed in these tumors (log_2_FC=1.7079, p=0.010), as IFNγ reduction activates Notch1 signaling and GSC marker CD133 expression (Jorgovanovic et al., 2020). Notably, epithelial-mesenchymal transition (EMT)-promoting genes downstream of TGFβ/PDGF, including direct ECM fibers (collagen type IV/VI, laminins, and actin) and ECM remodeling components (MMP9, syndecan-1, integrin receptors), were robustly upregulated in CD8a^lo^ specimens, further promoting stemness and invasiveness (Hao et al., 2019; Yu et al., 2021).

These contrasting CTL- vs. GSC-promoting transcriptional signatures which were jointly present in control GBMs relative to TAN-depleted GBMs aligned with historical models of antitumoral N1 and protumoral N2 neutrophil polarization states, wherein IFNγ induces the former phenotype and TGFβ the latter (Fridlender et al., 2009). Traditionally, inflammatory N1 neutrophils upregulate TNFα and ICAM1 expression, while N2 neutrophils promote invasion/angiogenesis and tumor stemness, analogous to the profiles we observed in CD8a^hi^ and CD8a^lo^ tumors, respectively (Anselmi et al., 2022). Broadly, our findings suggested that, in the presence of T-cells, an inflammatory positive feedback loop arises between hybrid TANs and IFNγ-producing T cells, whereas, in T cell-barren tumors, TAN-presented antigen cannot elicit a response and oncogenic signaling predominates.

### TANs in T-cell deficient mice promote GBM growth via GSC enrichment

To test the dependence of TAN-mediated tumor suppression on T cell co-infiltration, we evaluated the effect of αLy6G administration on constitutively T cell-deficient, athymic NuJ mice, which were implanted with identical BGL1 murine gliomas. Depletion was achieved in this model with long-term efficiency comparable to that seen in the BalbC strain (3 week efficiency: 48.7% systemically, 48.4% intratumorally; **Supp. Fig 3D**).

Contrary to our findings in immunocompetent mice, αLy6G-treatment attenuated tumor growth in the athymic model, confirming the necessity of T cells for anti-tumoral TAN function (**Fig. 3H**). Further, while neutrophil depletion had no effect on overall immune (CD45^+^) or myeloid (CD11b^+^) infiltration in both the BalbC and NuJ models (**Supp. Fig. 3E**), neutrophil depletion markedly diminished MHCII expression only in the latter, corroborating the aforementioned importance of T cell-associated IFNγ signaling on antigen presentation (**Supp. Fig. 3F**). In fact, the extent of MHCII^+^ neutrophil abundance corresponded to CD8^+^ T-cell levels across models (P<0.001; **Fig. 3I**), again underscoring the reciprocal nature of TAN-T cell crosstalk in stimulating TAN APC features and effecting T-cell-mediated cytotoxicity (Zhao et al., 2022).

To determine whether neutrophil enhancement of GBM growth in the absence of T cells was attributable to the predominance of PDGF/HIF1α signaling enriching GSCs over IFN signaling, we evaluated tumoral expression of the GSC markers CD133 and SOX2. While TANs in immunocompetent mice did not induce stemness, CD133 and SOX2 expression were increased by 57.6% (6.1% vs 14.3%, p=0.020) and 44.6% (14.8% vs 26.8%, p=0.024), respectively, in neutrophil-competent NuJ mice compared to αLy6G-treated NuJ mice (**Fig. 3J**). To then evaluate if TANs in isolation promoted stemness beyond surface marker upregulation, we cultured GBM43-derived GSCs in the presence of control or patient TAN-conditioned media (TANcm), finding that the latter did, indeed, yield larger GBM43-derived neurospheres (P<0.001; **Supp. Fig. 3G**).

### Immunostimulatory and GSC-promoting phenotypes reside jointly in a single noncanonical TAN polarization state distinct from conventional cytotoxic TANs

Our observation that TANs could both enrich GBM stemness and simultaneously immunologically attenuate GBM growth prompted us to interrogate whether these contrasting phenotypes were purely attributable to T cell colocalization, or instead reflected TAN heterogeneity. To that end, we employed scRNA-seq on patient-matched TANs and PBNs from newly diagnosed GBM. After exclusion of non-viable and contaminant immune cells (CD3D/CD3E/CD4+ T cells, CD79A/B+ B cells, and CD14+ monocytes), 7463 PBNs and 11831 TANs were included in the final analysis.

Broadly, TANs and PBNs exhibited distinct and stratified clustering in UMAP space (**Fig. 4A; Supp. Fig. 4A**), recapitulating their contrasting phenotypes. Module scores were computed for genes defining the MHC class I and class II pathways, and consistent with our prior findings, TANs upregulated both modes of antigen presentation (Matsushima et al., 2013; Singhal et al., 2016). In particular, the magnitude of this upregulation was greater for class II presentation (t_MHCII_=121.11, p<0.001; t_MHCI_=25.71, p<0.001), driven largely by the MHCII subunit HLA-DRA (log_2_FC=2.98), associated invariant chain CD74 (log_2_FC=3.02), and activation marker CD83 (log_2_FC=1.46). Further corroborating this immunostimulatory signature, the top 50 genes upregulated by TANs included cytokines IL1B (log_2_FC=3.82) and CXCL8 (log_2_FC=1.65), as well as macrophage inflammatory protein-1 (MIP-1) subcomponents CCL4 (log_2_FC=3.98) and CCL3 (log_2_FC=2.98; **Fig. 4B; Supp. Table 5**). Many of these genes are conventionally expressed by dendritic cells (CD83) and macrophages (MIP-1, IL1B), again illustrating a hybrid phenotype with mixed myeloid properties. However, we found that these cells still robustly expressed S100A8 and S100A9 (95.5% and 92.4% of TANs, respectively), which together encode the predominant intracellular protein in neutrophils, calprotectin. In fact, expression of these genes was significantly greater among TANs than PBNs (S100A8 log_2_FC=0.78, S100A9 log_2_FC=0.91), suggesting activation at a transcriptional level consistent with our previous morphologic findings (Sprenkeler et al., 2022).

**Figure 4.**
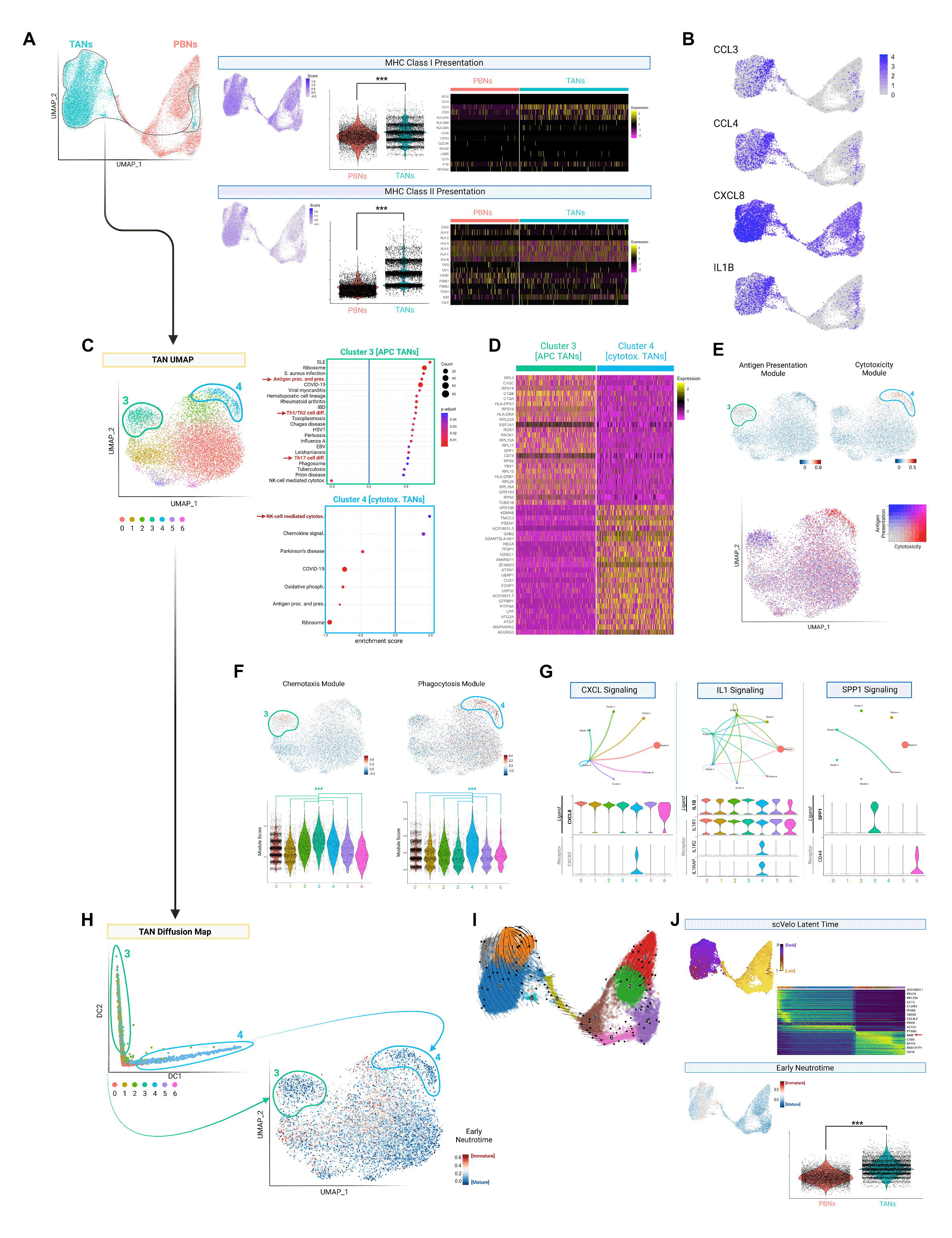
Single-cell RNA-sequencing demonstrates TAN polarization into canonical and hybrid subsets from precursors not seen in circulation. **(A)** *Left:* scRNA-seq of patient-matched PBNs and TANs, jointly represented in UMAP space. *Right:* Module scores for MHC class I and class II-related gene expression, visualized per cell (feature plot) and compared between neutrophil sources (violin plot). Heat maps depicting constituent genes shown. **(B)** Feature plot depicting expression of select cytokines. **(C)** UMAP visualization of TANs in isolation, with KEGG pathway enrichment analysis of clusters 3 (green, n=866 or 7.3%) and 4 (blue, n=488 or 4.1%). Only significantly up-or down-regulated pathways shown; select pathways highlighted in red. **(D)** Heatmap depicting the top 50 differentially expressed genes between clusters 3 (APC TANs) and 4 (cytotoxic TANs). **(E)** Feature plots depicting module scores calculated using constituent genes from the “Antigen Processing and Presentation” and “NK-cell mediated cytotoxicity” KEGG pathways (upper plots). Combined representation shown below. **(F)** Feature and violin plots depicting module scores for chemotaxis-and phagocytosis-related genes. Only comparisons with clusters 3 and 4 are shown. **(G)** Network graphs from CellChat ligand-receptor analysis of TANs, with dot size representing cluster size and connecting arrows colored to match the ligand-expressing cluster. Matched violin plots shown below. **(H)** Diffusion map revisualization of TANs, demonstrating clusters 3 and 4 (circled) to localize to opposite ends of the resulting continuous distribution. Corresponding feature plot of an ‘Early Neutrotime’ module in the original TAN UMAP, which is computed based on genes associated with neutrophil immaturity, and thus lower (bluer) scores indicate greater maturity. **(I)** RNA velocity vectors (scVelo) illustrating developmental relationships between TAN and PBN clusters in the combined dataset. **(J)** Quantification of neutrophil maturity by scVelo latent time (*above*) and Early Neutrotime (*below*, inversely proportional to maturity), with scores depicted as feature plots. For latent time, heatmap of top transitionally expressed genes shown. For Early Neutrotime, violin plot comparing PBNs and TANs shown. *Data are represented as means ± SD*. *\*p<0.05, **p<0.01, ***p<0.001*

In discerning whether these hybrid properties identified TANs wholly or a focal subset, we observed that of six TAN clusters, cluster 3 (866 cells=7.3% of TANs) enriched genes in the KEGG pathway corresponding to antigen presentation/processing and Th1/2/17 differentiation (**Fig. 4C-D; Supp. Fig. 4B; Supp. Table 6**). In contrast, cluster 4 (488 cells=4.1% of TANs) displayed a more conventional phenotype, upregulating NK-cell-mediated cytotoxicity genes (**Supp. Table 7**). Combined representation of module scores corresponding to these pathways further highlighted the stark dichotomy between these APC (cluster 3) and cytotoxic (cluster 4) TAN clusters (**Fig. 4E**). Additionally, the former population exhibited a chemotactic profile characterized by CCL3/CCL4/C1q/IL1 expression while the latter robustly expressed motility and phagocytosis-related genes (**Fig. 4F; Supp. Fig. 4C**), suggesting contrasting primary effector mechanisms: leukocyte recruitment and activation by APC TANs, and canonical microbicidal function by cytotoxic TANs. In fact, when frescoed against the integrated UMAP space, cytotoxic TANs clustered among PBNs (**Supp. Fig. 4D**), representative of their classical PMN-like transcriptional features. Similarly, 41 of the top 50 differentially expressed genes defining cytotoxic TANs relative to other TAN populations were identical to genes upregulated by PBNs relative to TANs (**Supp. Fig. 4E**).

When we applied the ligand-receptor prediction algorithm, CellChat, to discern if APC and cytotoxic TANs interacted, we found that cytotoxic TANs displayed a conventional responder phenotype, upregulating receptors for cytokines (CXCL8, IL1) expressed by other TANs (**Fig. 4G**). Meanwhile, in addition to these cytokines, APC TANs also robustly expressed *SPP1* (osteopontin). As osteopontin is a known promoter of GSC enrichment (Pietras et al., 2014), we tested whether it was also responsible for mediating the TAN-induced GBM stemness we had observed both in T cell-deficient murine hosts and *in vitro*. Indeed, we observed via ELISA that, across patients, this protein was secreted in greater quantity by TANs than PBNs (**Supp. Fig. 4F**), and that OPN blockade impeded the stimulatory effect of TANcm on expression of stemness genes *Nanog* and *Oct4* (P<0.01; **Supp. Fig. 4G**) and GSC neurosphere formation (P<0.05; **Supp. Fig. 4H**). Critically, the observation that noncanonical TANs exhibited both inflammatory antitumoral and stem-promoting protumoral features reinforced the centrality of T cell infiltration in determining the net effect of TANs on tumor biology.

### Trajectory and pseudotime analysis reveal that hybrid TAN development occurs intratumorally from immature precursors

Given the stark differences between cytotoxic and hybrid TANs, we next applied diffusion mapping, an alternative clustering approach optimized to explore differentiation trajectories, to delineate if these two phenotypes represented alternative polarization programs. Indeed, we observed that cells corresponding to clusters 3 and 4 localized to opposite ends of the resulting continuous distribution. The distinct transcriptional profiles of clusters 3 and 4 and their divergence from PBNs implied the origin of these clusters from distinct local progenitors (**Fig. 4H**). Moreover, while cytotoxic TANs closely resembled PBNs exposed to the tumor secretome, they notably upregulated genes (e.g., OSM, HLA-DRA) that could *not* be induced in culture (**Supp. Fig. 4I; Supp. Table 6**), suggesting the presence of intratumoral precursors that are less differentiated and more plastic than circulating PBNs.

To query TANs for this immature subset, we calculated Early Neutrotime scores based on a recently published transcriptional module tracing neutrophil development (Grieshaber-Bouyer et al., 2021). While cytotoxic and hybrid TANs were relatively differentiated, most TANs displayed high Early Neutrotime scores consistent with developmental naivety. The corresponding implication – that hybrid TANs arose intratumorally rather than peripherally – was further validated by an alternative, unsupervised pipeline (scVelo), in which no RNA velocity vectors directed differentiation from PBNs towards TANs (**Fig. 4I**).

Both early granulopoietic markers (Neutrotime) and transitional expression patterns (scVelo) demonstrated that immature neutrophils were exclusive to the TME, as PBNs universally exhibited low Early Neutrotime and high latent time scores (**Fig. 4J; Supp. Fig. 4J**). Of note, the top dynamically expressed genes (n=264) driving the latter index included canonical neutrophil maturity markers (MME, PTPRC; **Supp. Table 7**), suggesting that this disparity in latent time reflected maturity. In general, not only did PBNs upregulate additional markers of terminal differentiation (CXCR2, log_2_FC=0.62; ITGAM, log_2_FC=0.53), but they also resembled mature resting PMNs phenotypically, robustly expressing many tertiary granule, phagocytosis, and respiratory burst genes (**Supp. Fig. 4K**). In contrast to the chemotactic cytokines robustly expressed by TANs, PBNs had greater expression of genes pertaining to mechanical motility, including the MYO1F/ADAM8 cytoskeletal complex (log_2_FC=1.71, 0.45), cytoskeletal reorganization element DOCK2 (log_2_FC=1.30), and transepithelial migration mediating JAML (log_2_FC=0.69). Finally, unlike the inflammatory signature identified in hybrid TANs, PBNs displayed an immunoregulatory signature prototypical of resting PMNs, upregulating ENTPD1 [CD39] (log_2_FC=1.99), GIT2 (log_2_FC=1.83), and SEMA4D (log_2_FC=1.21), which generate immunosuppressive adenosine, attenuate TLR signaling, and suppress premature activation, respectively (Minns et al., 2021; Wang and Chen, 2018; Wei et al., 2014).

### Induction of the hybrid phenotype occurs exclusively in immature murine neutrophils

As our scRNA-seq data suggested that the differentiation of hybrid TANs occurred entirely within the TME from immature progenitors, we first validated that PBNs could not be directed to express MHCII. Indeed, when we exposed patient and healthy control PBNs to U251 tumor-conditioned media or cocultured them with U251 cells outright, we were unable to induce MHCII expression (P=0.1-0.2; **Fig 5A**). We subsequently explored the possible roles of self-recognition and other immune cells in this phenomenon, coculturing patient PBNs with matched tumor cells, either with infiltrating leukocytes removed or present. Again, we failed to observe surface-level MHCII expression (P=0.2-0.9; **Fig. 5A**).

**Figure 5.**
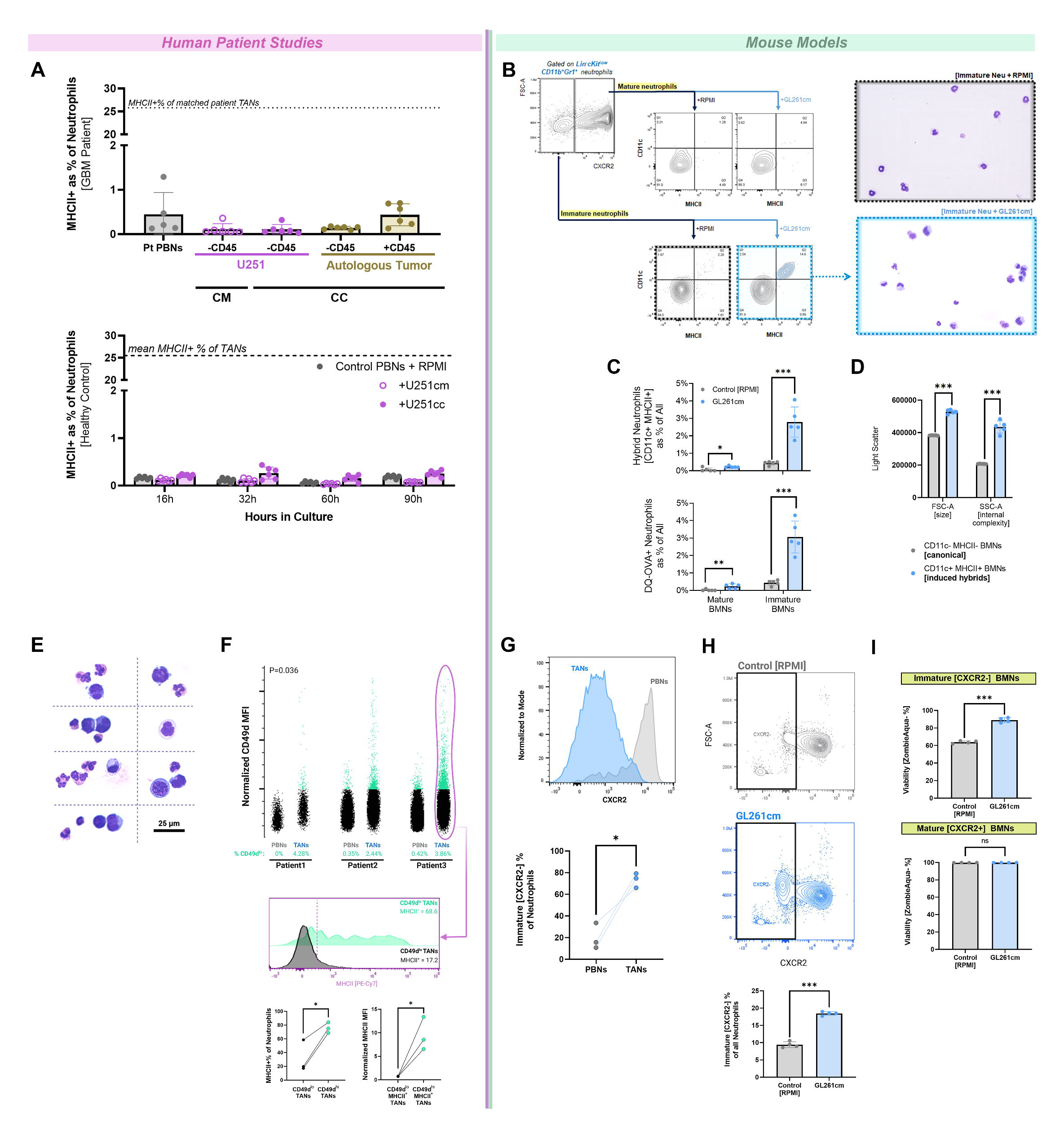
MHCII^+^ TANs with hybrid features arise from immature neutrophils rather than PBNs. **(A)** Surface MHCII expression in patient (*above*; n=6/group) and healthy volunteer (*below;* n=6/group) PBNs exposed to various inducing stimuli. PBNs were cultured in U251cm or directly with U251 cells (coculture, “U251cc”) for 16h, and for incremental intervals thereafter in the healthy volunteer arm. Patient PBNs were also cultured with autologous tumor, with or without CD45 immune cells present. **(B)** Representative contour plots (*left*) of MHCII and CD11c expression by FACS-purified mature (CXCR2^+^Ly6G^hi^) and immature (CXCR2^-^Ly6G^lo/hi^) Lin^-^ CD11b^+^Gr1^+^ BMNs cultured in control or GL261-conditioned media for 48h. Wright-Giemsa stained cytospins of immature BMNs (*right*) depict hybrid features (dendritic processes, irregular nuclei) in GL261cm-exposed cells. Images of induced hybrids are stitched together in the bottom right picture for clarity. **(C)** Quantification of APC feature induction in immature and mature BMNs exposed to GL261cm, including hybrid marker expression (P_mature_=0.01, P_immature_<0.001) and DQ-OVA uptake/processing (P_mature_=0.008, P_immature_<0.001). Induction of both attributes was greater among immature BMNs (F_hybrid_=30.84, P<0.001; F_DQ-OVA_=32.51, P<0.001). N=5/group. **(D)** Size (FSC) and internal complexity (SSC) of induced hybrid BMNs compared to CD11c^-^MHCII^-^ canonical BMNs. N=5. **(E)** Myeloblastic cells observed in Wright-Giemsa stained cytospins of patient TANs, with non-segmented nuclei, azurophilic hue, and a generally enlarged profile. **(F)** *Above:* Dot plots of immaturity (CD49d expression) in 3 paired PBN-TAN samples. *Below:* Representative histogram of MHCII expression in CD49d^hi^ vs CD49d^lo^ TANs, with quantification across paired samples. **(G)** Representative histogram of TAN and PBN maturity (CXCR2 expression) in GL261 GBM-bearing C57BL/6J mice, with quantification below (n=3, P=0.036). **(H)** Representative contour plots of mixed bone marrow suspensions from C57BL/6J mice cultured in control or GL261-conditioned media for 72h, with quantification below (n=3/group). **(I)** Effect of GL261cm on BMN viability, assessed by ZombieAqua staining, in C57BL/6J BMNs (n=4/group). *Data are represented as means ± SD*. *\*p<0.05, **p<0.01, ***p<0.001*

Next, to explore the inducibility of more developmentally plastic cells – analogous to those observed in our scRNA-seq analysis – we leveraged murine bone marrow neutrophils (BMNs). As with human PBNs, exposure of murine C57BL/6J PBNs isolated by negative magnetic selection to syngeneic GL261 tumor-conditioned media (GL261cm) did not yield MHCII expression after 72h (P=0.5; **Supp. Fig. 5A**). However, *ex vivo* cultures of FACS-purified mature Lin^-^CD11b^+^Gr1^+^**CXCR2^+^Ly6G^hi^** and immature Lin^-^CD11b^+^Gr1^+^**CXCR2^-^Ly6G^lo/hi^** BMNs demonstrated that the latter could *robustly* be induced to express MHCII in response to tumor-secreted proteins (**Fig. 5B; Supp. Fig. 5B**) (Evrard et al., 2018; Grieshaber-Bouyer et al., 2021; Xie et al., 2020). Moreover, the resultant MHCII^+^ population co-expressed the dendritic cell marker CD11c (P=0.0003 immature, P=0.01 mature; **Fig. 5C upper panel**) and possessed the functional capacity to process exogenous peptide antigen (P=0.0002 immature, P=0.008 mature; **Fig. 5C lower panel; Supp. Fig. 5C**), again reflecting a hybrid phenotype (Matsushima et al., 2013). Indeed, conditioned immature BMNs developed oval nuclei and dendritic projections, while non-conditioned immature BMNs maintained classical neutrophil morphology with ringed central nuclei and smooth cellular membranes. Quantitatively, these induced hybrids were larger and more structurally complex, as proxied by FSC (P<0.001) and SSC (P<0.001), respectively (**Fig. 5D**). All findings were confirmed in Balb/cJ mice using an alternative syngeneic glioblastoma model (BGL1 in Balb/cJ mice; P<0.001; **Supp. Fig. 5D**).

### Histologic and cytometric identification of immature TANs in patient and murine GBMs

The exclusive inducibility of immature neutrophils to achieve a hybrid TAN phenotype, together with our scRNA-seq data, suggested that precursor immature TANs must infiltrate the GBM TME at a cellular level. Indeed, though we generally found patient TANs to be hypersegmented, our analysis of Wright Giemsa-stained cytospins revealed a noticeable subset of azurophilic cells with high nuclear-to-cytoplasmic ratios among TANs, resembling early lineage myeloblast-like cells (**Fig. 5E**); these cells constituted 1.10% of imaged TANs (n=1824 cells), but were entirely absent in circulation. Subsequent flow cytometric analysis of neutrophil maturity in 3 paired samples of patient TANs and PBNs yielded consistent results, demonstrating that CD49d^hi^ pre-neutrophils (preNeus) were highly enriched in the tumor, comprising 3.52% [95% CI: ±2.40%] of TANs and only 0.26% [95% CI: ±0.56%] of PBNs (p=0.036; **Fig. 5F**).

Collectively, these findings indicated that early lineage neutrophils accumulated in the human GBM TME. Moreover, we observed that among these CD49d^hi^ TANs, hybrid neutrophils were 3.07 times (p=0.042) more common and had 12.33 times (p=0.0495) greater average surface MHCII expression than among CD49d^lo^ TANs, recapitulating the link between developmental immaturity and differentiation into the hybrid phenotype. Critically, as CD49d (integrin α_4_) also serves as a functional receptor for osteopontin, which is in turn secreted by hybrid TANs, this suggested that hybrid TANs may cyclically promote their own expansion by augmenting precursor recruitment (Cao et al., 2019). In fact, noncanonical TANs were also found to robustly upregulate C1q, which is similarly involved in HSPC homing (Jalili et al., 2010).

To evaluate if this intratumoral accumulation of immature neutrophils was preserved in syngeneic murine models of GBM, we characterized the relative contribution of immature CXCR2^-^Ly6G^lo/hi^ and mature CXCR2^+^Ly6G^hi^ TANs in GL261 tumor-bearing C57BL/6J mice. Consistent with the finding that preNeus infiltrated human glioblastoma tumor specimens, immature neutrophils were disproportionately enriched in murine tumor samples compared to blood (P<0.05; **Fig. 5G**). Moreover, the tumor secretome preserved viable neutrophils in this relatively dedifferentiated state; in mixed bone marrow cultures, 18.5% [95% CI: ±0.9%] of neutrophils remained in the immature state after 48h when exposed to GL261 tumor-conditioned media (GL261cm), compared to 9.4% [95% CI: ±1.4%] in control conditions (p<0.0001; **Fig. 5H**). Further, GL261cm retarded early apoptosis specifically in these immature neutrophils (P<0.001; **Fig. 5I**).

### Immature TANs infiltrate GBM from adjacent skull bone marrow

We then sought to determine if the GBM microenvironment was specifically chemotactic for these immature, relatively plastic neutrophils. Mixed C57BL/6J bone marrow isolates were seeded atop transwell inserts and allowed to migrate towards either control media or GL261 tumor-conditioned media for 16 hours; flow cytometric characterization of migrated cells demonstrated enrichment in both immature neutrophils and overall granulocyte-monocyte precursors (GMPs) in the conditioned media group (**Fig. 6A;** p_GMP_=0.005, p_immNeu_=0.003).

**Figure 6.**
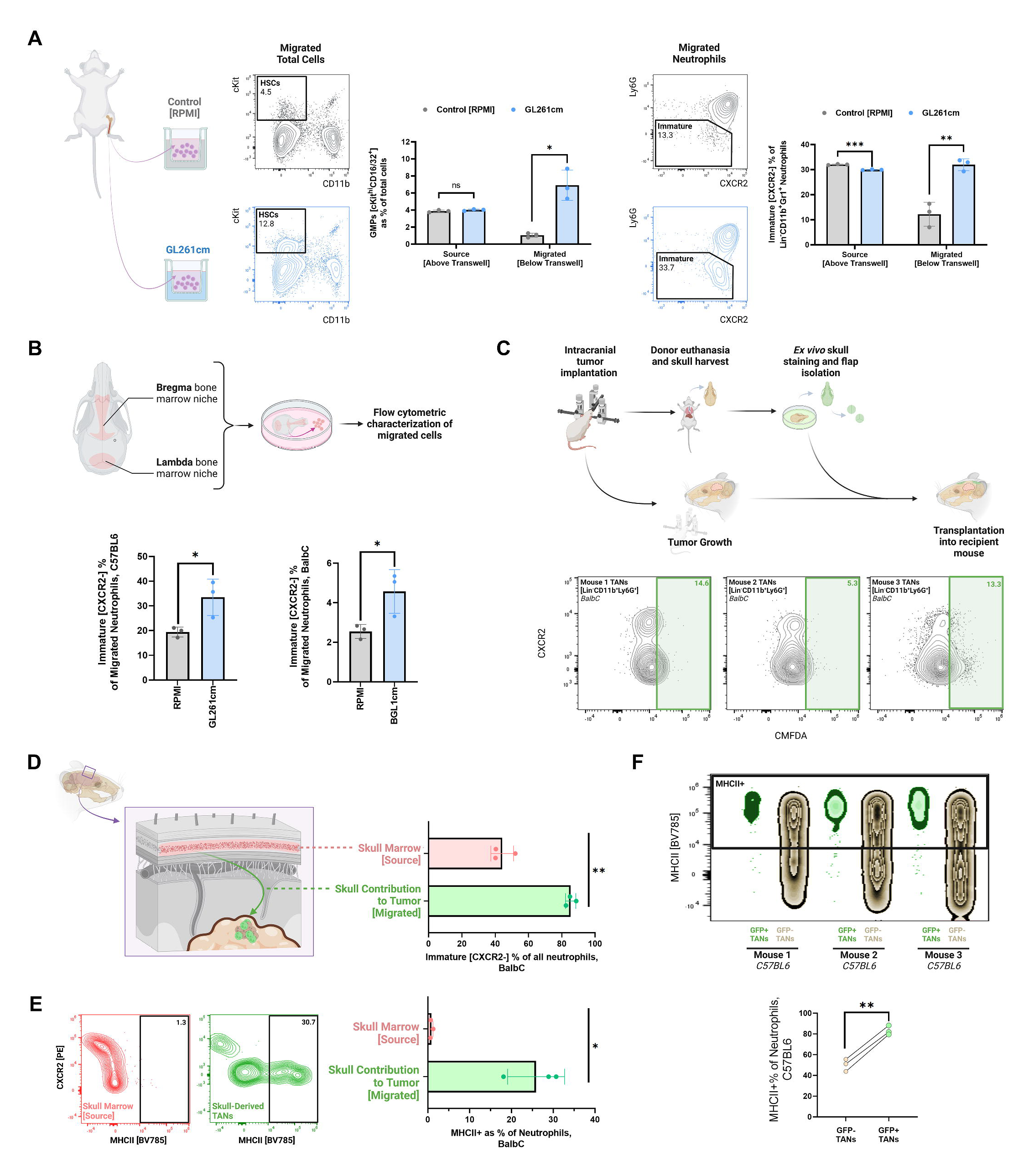
GBM recruits immature neutrophils from local skull marrow *in vivo* and subsequently induces MHCII expression. **(A)** 16h transwell migration of mixed C57BL/6J bone marrow isolates towards control or GL261-conditioned media (n=3/group), with unmigrated (above filter, “source”) and migrated (below filter) cells quantified by flow cytometry for cKit^hi^CD11b^-^ CD16/32^+^ GMPs (P_source_=0.232, P_migrated_=0.027) and immature neutrophils (P_source_<0.001, P_migrated_=0.008). Experimental schematic and representative contour plots of HSC (Lin^-^cKit^hi^CD11b^-^) and immature neutrophil (CXCR2^-^) gating shown. **(B)** *Above:* Schematic of murine calvarial bone marrow niches and *ex vivo* assay. *Below:* Quantification of immature neutrophil migration from calvaria cultured in control or syngeneic tumor-conditioned media for 48h (n=3/group). P_C57BL/6J_=0.045, P_BalbC/J_=0.043. **(C-E)** BGL1-bearing BalbC/J mice (n=3) were transplanted with CMFDA-stained skull flaps, and tumors were harvested after 72h for analysis by flow cytometry. **C:** *Above:* Schematic of skull flap staining and transplantation protocol. *Below:* Contour plots of TANs from each mouse, with CMFDA^+^ skull-derived cells gated based on matched FMOs. Values indicate % of all TANs. **D-E:** Immature (D) and MHCII+ (E) proportion of neutrophils contributed from the skull marrow to the tumor, compared to those in the skull itself (“source”). N=3/group, P_immature_=0.003, P_MHCII_=0.024. Representative contour plots shown in E. **(F)** Zebra plot of MHCII expression by definitively skull-derived (GFP+) and other (GFP-) TANs in 3 GL261-bearing C57BL/6J mice, which were transplanted with UBC-GFP skull flaps prior to tumor inoculation. Quantification below, P=0.002. *Data are represented as means ± SD*. *\*p<0.05, **p<0.01, ***p<0.001*

Then, in light of a previous study demonstrating that the inflamed brain after stroke has a high propensity to recruit neutrophils from local skull bone marrow rather than homogenously from all marrow sites in the body (Herisson et al., 2018), we investigated whether this myeloid reservoir contributed immature neutrophils to the GBM microenvironment. Calvaria including bregma and lambda were harvested from healthy C57BL/6J and Balb/CJ mice, then cultured in control media (RPMI) or tumor-conditioned media (GL261 and BGL1, respectively) for 48h (**Fig. 6B**). Consistent with our transwell findings, emigrated neutrophils in the presence of conditioned media were disproportionately immature (C57BL/6J: 33.5% vs 19.4%, p=0.045; Balb/CJ: 4.6% vs 2.5%, p=0.043). Subsequent *in vivo* characterization of BMNs in healthy and GL261 tumor-bearing C57BL/6J mice demonstrated that the skulls of tumor-bearing mice had fewer immature neutrophils than healthy skulls (38.2% vs 45.9%, p=0.048; **Supp. Fig. 6A**), further suggesting extensive tumoral recruitment of these cells from local marrow. By contrast, the composition of systemic (femur) BMNs was unaffected by tumor implantation.

To definitively demonstrate neutrophil chemotaxis from skull to tumor, dual bregma/lambda skull bone flaps were harvested from euthanized donor mice, labeled *ex vivo* with a CMFDA green fluorescent dye, and transplanted onto syngeneic BGL1 tumor-bearing Balb/cJ mice for 72h (**Supp. Figs. 6B-C**). Flow cytometric characterization of the resultant tumors universally (n=3) demonstrated infiltration by fluorescent, and thus skull-derived, neutrophils (**Fig. 6C**). These skull-derived TANs (S- TANs) were more immature in composition than the overlying skull bone marrow in non-transplanted tumor-bearing mice (85.4% vs 44.2%, p=0.003; **Fig. 6D**), again suggesting their preferential recruitment and chemotaxis to the microenvironment. Moreover, while BMNs in the overlying skull marrow did not appreciably express MHCII, S-TANs robustly expressed this marker (0.9% vs 25.9%, p=0.02; **Fig. 6E**), confirming that these cells retained their phenotypic plasticity. This observation reinforced a chronological sequence wherein skull BMNs first transit to the GBM TME, then develop the hybrid phenotype.

Given the abundance of immature neutrophils among S-TANs, we anticipated enhanced MHCII induction within this population compared to the remaining TANs. Indeed, hybrid TANs were more frequent among S-TANs than among other TANs (18.8% vs 25.9%, p=0.01; **Supp. Fig. 6D**), but this disparity was slight, likely secondary to the limited 72h interval between skull flap transplantation and tumor harvest, which was required given the transient nature of CMFDA staining.

Thus, to better longitudinally assess the phenotype of S-TANs, GFP-expressing skull bone marrow flaps from UBC-GFP mice were used instead. GFP+ skull flaps were transplanted onto wild-type C57BL6 mice immediately *prior* to GL261 tumor implantation, and tissue was harvested at 3 weeks. Though the transplanted flaps had limited viability and were extensively repopulated by GFP- bone marrow cells from the adjacent skull at endpoint (**Supp. Fig. 6E**), we still observed infiltration by GFP+ neutrophils in all mice (n=3; **Supp. Fig. 6F**). These S-TANs were again disproportionately immature compared to skull bone marrow in non-transplanted tumor-bearing mice (80.0% vs 36.7%, p=0.005). Critically, almost all (83.2%) S-TANs were MHCII+, whereas only half (50.2%) of the remaining TANs were MHCII+ (p=0.002; **Fig. 6F**). This suggested that, over the course of GBM growth, infiltrating skull marrow-derived neutrophils are fated to become long-lived hybrid TANs, and thus the skull marrow contributes potently to this population.

Finally, we observed that in addition to hybrid-fated neutrophils, the skull also contributed inflammatory macrophages [Lin^-^CD11b^+^Gr1^+^CD115^+^F4/80^+^] to the TME in both experimental models: Balb/cJ with CMFDA-stained transplants, and C57BL/6J with GFP+ transplants (**Supp. Fig. 6G**). Similar to S-TANs, these skull-derived macrophages (S-TAMs) were also polarized to an immunostimulatory state; compared to other TAMs, S-TAMs were enriched in MHCII^hi^ M1-like cells (Balb/cJ: 7.7% vs 21.5%, p=0.036; C57BL/6J: 9.4% vs 30.6%, p=0.019). Thus, our findings implicated the skull bone marrow as a potential driver of CD4 T cell responses through its multimodal supply of antigen-presenting myeloid cells to the GBM TME.

### Selective skull marrow ablation impedes hybrid TAN infiltration and accelerates GBM growth

To determine whether these skull-derived immune cells bore meaningful antitumoral relevance, as suggested by their inflammatory polarization, we selectively ablated the skull marrow through targeted cranial irradiation of body-shielded C57BL/6J mice 24h prior to syngeneic GL261 glioma implantation. We validated radiation-induced depletion of skull marrow immune cells, its durability over the experimental course, and its lack of off-target effects on native brain immune populations, observing minimal skull marrow immune repopulation (average: 8.6% repopulation per week; **Supp. Fig. 7A**) and comparable microglial infiltration to non-irradiated controls (P=0.7; **Supp. Fig. 7B**). Conversely, to determine whether promoting the egress of skull marrow cells into GBM could be a therapeutic strategy capable of attenuating GBM growth and prolonging survival, an additional cohort of non-irradiated mice were treated 9 days after GBM implantation with a single intracalvarial dose of the CXCR4 antagonist AMD3100, which has been reported to induce the egress of myeloid cells from the skull marrow into the meninges (Cugurra et al., 2021).

Compared to controls, median overall survival was 22.7% shorter in skull-irradiated mice (P<0.001) and 20.5% longer in AMD3100-treated mice (P=0.006), suggesting that altering the skull marrow-mediated antitumoral response had direct effects on the survival of GBM-bearing mice (**Fig. 7A**). Accordingly, tumor growth was accelerated in skull-irradiated mice and attenuated in the AMD3100-treated group, with skull-irradiated tumors measuring 10.8-fold larger by BLI at day 15 than controls (P<0.001), which were in turn 7.7-fold larger than AMD3100-treated tumors (P=0.002) (**Fig. 7A**).

**Figure 7.**
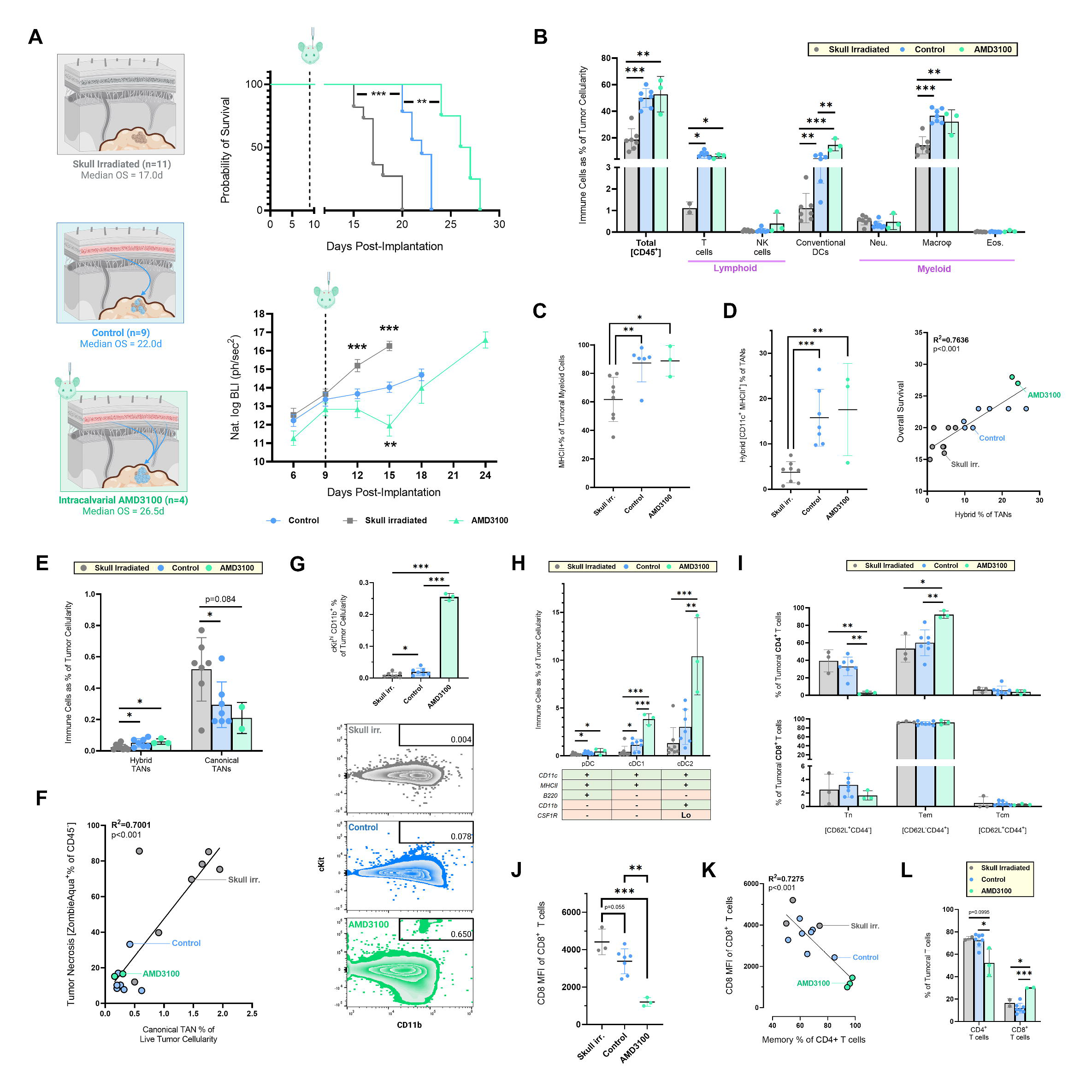
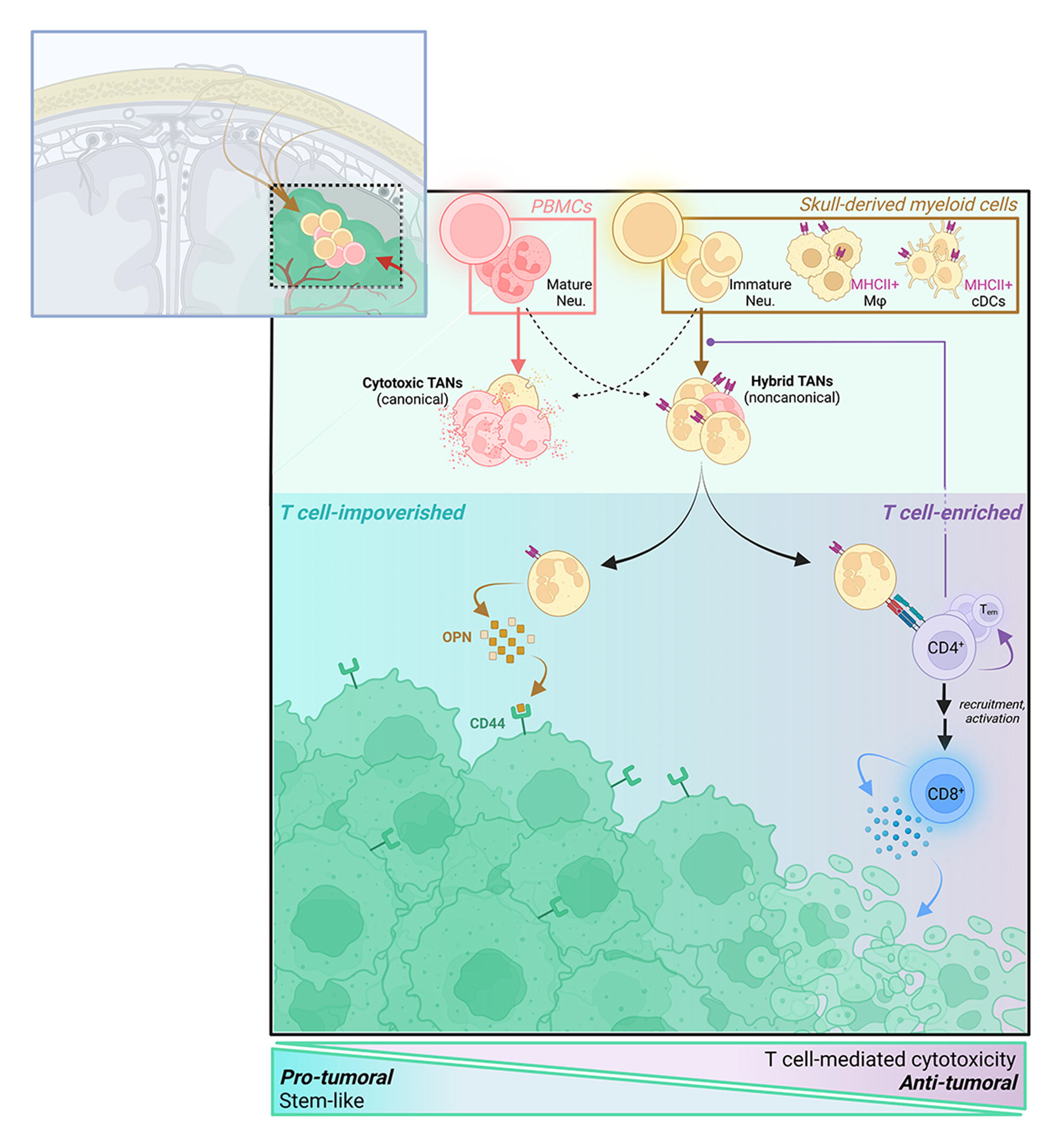
Altering the release of skull marrow precursors impacts the tumor immune profile and survival of GBM-bearing mice. **(A)** GL261-bearing C57BL/6J mice received either 13 Gy skull irradiation (n=11) pre-inoculation or 10 µg of intracalvarial AMD3100 at POD9 (n=4; dashed line). Kaplan-Meier survival (P_irrad_<0.001, P_AMD3100_=0.006) and tumor growth by BLI (P_irrad_<0.001 at POD15-18, P_AMD3100_=0.002) shown, with all comparisons relative to controls (n=9). **(B)** Flow cytometric quantification of immune populations as a proportion of endpoint tumor bulk. Total immune cells (P_control_<0.001, P_AMD3100_=0.001), T cells (P_control_=0.019, P_AMD3100_=0.028), cDCs (P_control_=0.002, P_AMD3100_<0.001), and macrophages (P_control_<0.001, P_AMD3100_=0.007) were decreased in skull-irradiated mice. cDCs were further enriched in AMD3100-treated mice compared to controls (P=0.001). **(C)** MHCII expression among tumor-infiltrating myeloid cells, enriched in control (P=0.003) and AMD3100-treated mice (P=0.023) compared to skull-irradiated. **(D)** *Left:* Hybrid polarization of TANs, greater in controls (P<0.001) and AMD3100 (P=0.004) than skull-irradiated. *Right:* Linear regression of overall survival on hybrid TAN polarization. **(E)** Contribution of neutrophil subsets to endpoint tumor bulk, with hybrid contribution reduced in skull-irradiated mice (n=8) (P_control_=0.025; P_AMD3100_=0.017) and canonical contribution conversely enriched (P_control_=0.034, P_AMD3100_=0.084). **(F)** Linear regression of endpoint tumor necrosis on canonical TAN infiltration. **(G)** Quantification and representative contour plots of myeloid precursor (CD11b^lo/hi^cKit^hi^) infiltration of endpoint tumors. P_irrad vs control_=0.048, all other P<0.001. **(H)** DC subsets, with marker definitions, in endpoint tumors. Enriched AMD3100- treated tumor infiltration by cDC1 and cDC2 compared to skull-irradiated (P<0.001) and controls (P_cDC1_<0.001, P_cDC2_=0.003), and by pDC compared to skull-irradiated (P=0.048). Infiltration by pDC (P=0.034) and cDC1 (P=0.035) also enriched in controls compared to skull-irradiated. **(I)** Memory T cell subsets (CD4 *above*, CD8 *below*) in endpoint tumors. Tn reduced (P_irrad_=0.008, P_control_=0.002) and Tem increased (P_irrad_=0.014, P_control_=0.007) in AMD3100-treated mice. **(J)** CD8^+^ T cell activation in endpoint tumors, proxied by a reduction in CD8 MFI, enhanced in AMD3100 tumors (P_control_=0.002, P_irrad_<0.001). **(K)** Linear regression of CD8 MFI (inversely related to activation) on memory polarization of CD4^+^ T cells. **(L)** Distribution of T cells, with fewer CD4^+^ (P=0.006) in AMD3100-treated tumors than in controls, and greater CD8^+^ than in controls (P<0.001) or skull-irradiated (P=0.040). *Data are represented as means ± SD. N=3-8 per group in all endpoint tumor analyses. For linear regressions, dots colored by experimental group*. *\*p<0.05, **p<0.01, ***p<0.001*

To delineate the relevant cell populations underlying these differences in survival, tumors were profiled by flow cytometry at endpoint. On average, overall immune infiltration as a proportion of tumor cellularity was 50.7% lower in skull-irradiated mice than controls (p=0.005; **Fig. 7B**), suggesting that the skull contribution to immune cells in the GBM TME was robust and that systemic marrow sites could not compensate for the loss of skull marrow. Specifically, this reduced immune infiltration in the GBM TME of skull-irradiated mice was attributable to a decrease in aggregate macrophage, dendritic (DC) and T cell infiltration, implicating calvarial marrow as a meaningful contributor to effective antigen presentation. Indeed, MHC-II expression among constituent myeloid cells was markedly reduced in skull-irradiated mice (P=0.007; **Fig. 7C**). Moreover, not only was there greater dendritic polarization of infiltrating TANs in control and AMD3100-treated mice than in skull-irradiated mice, but this trended linearly with overall survival across all three groups (R^2^=0.7636, p<0.001; **Fig. 7D**).

Conversely, skull-irradiated mice were so enriched in non-hybrid TANs that, despite their general paucity of immune cells, aggregate intratumoral infiltration by these non-hybrid neutrophils exceeded that observed in controls (P=0.034; **Fig. 7E**). In interrogating whether this skewed TAN profile favoring canonical TANs in skull-irradiated mice yielded a more cytotoxic tumor milieu, we observed that skull-irradiated tumors were indeed 4.5 times more necrotic than controls (60.7% vs. 13.5% dead cells, p<0.001), and that the extent of canonical TAN accumulation significantly correlated with this phenomenon across all three treatment groups (R^2^=0.7002, p<0.001; **Fig. 7F**). These data corroborated prior reports of TAN-mediated ferroptosis in GBM, suggesting that while skull-derived TANs (and other APCs) promote T cell recruitment/proliferation and a regulated adaptive immune response, an excess of systemically derived neutrophils in the absence of skull-derived TANs instead yields pathologic necrosis and a worse prognosis (Yee et al., 2020). In particular, neutrophil marker MPO has been reported as a key mediator of this protumorigenic response (Yee et al., 2020), which we found to be upregulated among PBNs compared to TANs (log_2_FC=9.5475, p<0.001) in our earlier Nanostring analysis of patient samples (**Supp. Table 2**).

### Intracalvarial administration of AMD3100 yields an influx of skull marrow-derived myeloid APCs and subsequent T-cell activation, slowing GBM growth

Not only did intracalvarial AMD3100 prolong overall survival in a manner that correlated with hybrid TAN polarization, but strikingly, intracalvarial AMD3100 temporarily reversed tumor growth following its administration, causing a 53.4% reduction in tumor size by BLI 6 days later (P<0.01; **Fig. 7A**). Notably, a sizeable population of CD11b^+^cKit^hi^ myeloid precursors was observed in AMD3100-treated GBMs that was only scantly present in control GBMs and even less so in skull-irradiated mice (P<0.001; **Fig. 7G**), indicating that intracalvarial AMD3100 was inducing release of hematopoietic precursors from the skull marrow and implicating this process in downstream tumor suppression.

We then sought to determine whether intracalvarial AMD3100 was impacting other differentiated cell populations through the release of these myeloid precursors from the skull marrow into GBM. While treatment did not alter intratumoral levels of most broad immune subsets (T cells, NK cells, neutrophils, eosinophils, macrophages) compared to controls (P>0.05; **Fig. 7B**), intratumoral DC infiltration was 3.3-fold higher in AMD3100-treated mice than in control mice (P=0.001; **Fig. 7B**). This disparity was not associated with an altered distribution of DC subsets, but rather, a uniform expansion of cDC1 (P<0.001) and cDC2 (P=0.003) cells (**Fig. 7H**). As classical DCs are generally thought to arise from rare circulating progenitors, their preponderance in this cohort suggested enhanced liberation of pre-cDCs from adjacent marrow (Merad et al., 2013; Naik et al., 2007; Sichien et al., 2017).

Finally, to ascertain whether this myeloid precursor and cDC influx manifested in downstream T cell stimulation, we interrogated effector and memory subsets among intratumoral CD8^+^ and CD4^+^ populations (Patente et al., 2018). Consistent with their widespread upregulation of MHC-II and relative abundance of cDC2 cells, AMD3100- treated tumors had greater CD4^+^ activation than controls, with a greater proportion of effector memory (T_em_) cells and a corresponding decrease in naïve (T_n_) cells (P_Tem_=0.007, P_Tn_=0.002; **Fig. 7I**). While this same phenomenon was not observed among CD8^+^ T cells, downregulation of the CD8 coreceptor – indicating early activation in advance of memory subset differentiation (Xiao et al., 2007) – was most prominent in AMD3100-treated mice, followed by controls and the skull-irradiated cohort (P<0.001; **Fig. 7J**). In fact, the extent of this downregulation directly correlated with the size of the CD4^+^ memory pool, suggesting the latter’s polarization into the CTL-stimulatory Th1 subtype (R^2^=0.7275, p<0.001; **Fig. 7K**). Indeed, CD8^+^ T cells as a whole were enriched in AMD3100-treated mice (**Fig. 7L**). Thus, intracalvarial AMD3100 prolonged survival of GBM-bearing mice by enhancing the anti-tumoral T-cell response through liberation of skull marrow myeloid precursors into GBM, where they differentiated into antigen-presenting MHC-II+ myeloid cells, including hybrid TANs and DCs.

## DISCUSSION

Tumors, which were once perceived as homogenous masses of cells, are now recognized as organ-like structures in which tumor cells engage in dynamic interactions with cells in their microenvironment. Of these interactions, those with TANs are among the least well understood. While early studies suggested that TANs were bystanders either being detected intravascularly as they circulate through the tumor or passively accumulating without actual function because it seemed unlikely that such short-lived cells could meaningfully modulate chronic and progressive diseases like cancer, our study refuted these assumptions in multiple ways. First, we found that TANs had an enhanced lifespan compared to PBNs. Second, we found that TANs, in fact, had a distinct PBN-independent lineage, arising instead from immature precursors in the skull marrow that arrive in GBM without traversing the circulation. Third, we found that GBM TANs function as APCs, driving an anti-tumoral immune response in the presence of T-cells.

APC-like hybrid neutrophils have been previously described in early-stage lung cancer (Singhal et al., 2016). That finding and our work build upon prior reports demonstrating that neutrophils can acquire an APC phenotype under non-cancerous inflammatory conditions (Ashtekar and Saha, 2003), evidenced by upregulation of MHC class II subunits and co-stimulatory molecules, such as CD80 and CD86 (Iking-Konert et al., 2005). Critically, we are the first to contextualize this property *in vivo*, made possible by our utilization of a syngeneic Balb/CJ model. Unlike prior studies of αLy6G-mediated neutrophil depletion, which were conducted using short depletion intervals administered well after glioma implantation in C57BL/6 mice (Alghamri et al., 2021; Chen et al., 2022; Fujita et al., 2010; Jeon et al., 2019; Magod et al., 2021; Wang et al., 2020), our choice of a murine strain with higher sensitivity to ablation (Boivin et al., 2020) enabled us to evaluate the gross effects of TANs longitudinally, highlighting the antitumoral role of early-infiltrative, hybrid-predisposed TANs (Magod et al., 2021; Singhal et al., 2016). In fact, polarization of TANs into APCs was indispensable to this tumor suppression, as when we eliminated hybrid precursors through skull irradiation, residual canonical TANs instead promoted prognostically unfavorable necrosis.

Our finding of these APC-like hybrid TANs has different implications in GBM patients than in other cancers because GBM is a notoriously T-cell depleted cancer. We found that, while neutrophil depletion accelerated tumor growth in immunocompetent mice, the opposite occurred in athymic mice. These findings are likely because, while the APC-phenotype of hybrid TANs in GBM predominates in the presence of T-cells, a stimulatory effect of these TANs on GSCs predominates in the absence of T-cells. Further work will be needed to determine how these findings translate in GBM patients who have reduced levels of circulating T-cells compared to healthy adults (Chongsathidkiet et al., 2018) and whose T-cells are functionally impaired (Woroniecka et al., 2018), but our demonstration that hybrid TANs can stimulate autologous T cell activation in an MHCII-dependent manner suggests that therapeutic strategies designed to enhance these interactions are worth exploring further.

Our work also includes the first ever scRNA-seq analysis of purified TANs, building upon a prior study of healthy volunteer circulating neutrophils (Xie et al., 2020) and studies in which scRNA-seq of entire tumors were integrated to gain insights into TAN heterogeneity (Salcher et al., 2022; Xue et al., 2022). Through multiple modifications of the standard processing protocol, we were able to generate abundant high-quality transcriptomic reads despite the fragility and low transcriptomic density of neutrophils, enabling the interrogation of intratumoral heterogeneity in a manner unobscured by interpatient variability. Our analysis demonstrated that while TANs ranged phenotypically from canonical (PBN-like) to APC-like, this polarization was not concordant with historical N1/N2 or classical/MDSC models, corroborating evolving efforts to replace the idea of an MDSC monolith with one that embraces myeloid plasticity (Hegde et al., 2021; Khan et al., 2020). Indeed, rather than individual immunosuppressive TAN clusters upregulating ARG1/iNOS (Veglia et al., 2021), we observed that dual protumoral and antitumoral properties existed within a single noncanonical population, the functional consequences of which depend integrally on the presence of T cells.

MDSC nomenclature notwithstanding, our scRNA-seq analysis did lend credence to the idea that immature myeloid cells accumulate intratumorally. Further, given our robust capture of diverse cell states, we were able to elaborate upon this concept, demonstrating through trajectory and maturity analyses that these precursors are not a static endpoint but instead give rise to other populations – including MHCII^+^ hybrid TANs. While the role of MHCII^+^ myeloid cells broadly in inducing T cell cytotoxicity has been established (Kilian et al., 2023), prior studies have dichotomized their origin into local and peripherally-derived. By contrast, our work defines skull marrow-derived cells as a third distinct lineage of intratumoral antigen presenting cells, not just in TANs but also in DCs and TAMs.

This finding represents an oncological perspective to an expanding body of literature surrounding brain border immunological surveillance, recently invigorated by the discovery that calvarial bone marrow supplies immune cells to the CNS through microscopic vascular channels crossing the inner table of the skull (Herisson et al., 2018). The initial identification of this phenomenon demonstrated trafficking of myeloid cells to brain tissue after stroke and meningitis (Herisson et al., 2018), monocytes and neutrophils to CNS parenchyma upon injury (Cugurra et al., 2021), and B cells to the meninges under homeostasis (Brioschi et al., 2021). Our work builds upon those findings by demonstrating that the skull marrow contributes myeloid cells to GBM capable of antigen presentation and T-cell activation. The greater ability of skull marrow-derived cells to elicit a functional cytolytic response against GBM compared to systemic bone marrow could reflect an ability of the calvarial-meningeal path of immune cell development to provide the CNS with a constant supply of immune cells educated by CNS antigens (Pulous et al., 2022).

These properties provided compelling evidence that mobilizing skull-derived progenitors could meaningfully suppress tumor growth, to which end we pursued intracalvarial administration of AMD3100. Our use of this strategy builds upon previous studies showing that the skull bone is permeable to small-molecular-weight compounds that can affect meningeal immune cells after brain injury (Roth et al., 2014) by demonstrating the ability of an intracalvarial small molecule to influence the immune cell makeup of GBM parenchyma. Importantly, while recent studies have leveraged this compound to explore skull marrow contribution to neuroinflammation, we present the first therapeutic application of AMD3100 in this context, justifying the continued testing of treatments that exploit this novel intracalvarial administration route. Should further studies validate this approach and demonstrate a significant skull contribution to antigen-presenting myeloid cells in patients, this strategy should be readily implementable clinically.

## ACKNOWLEDGEMENTS

The authors would like to thank Akane Yamamichi, MD, PhD and Pavlina Chuntova, PhD for their assistance with animal experimentation and flow cytometry panel design. M.K.A. was supported by the NIH (1R01CA227136, 2R01NS079697, and 1R01NS123808). This study was supported in part by HDFCCC Laboratory for Cell Analysis Shared Resource Facility through grants from NIH (P30CA082103 and S10 OD021818-01).

## AUTHOR CONTRIBUTIONS

M.L. and A.S.B. conceptualized the study, designed and performed experiments, and analyzed and interpreted data. S.S.S., G.Y., A.T.N., A.L., and D.S. performed *in vitro* experiments and microscopy. J.S.Y. and S.G. performed *in vivo* experiments. J.J. and H.B. processed patient tissue and generated single-cell RNA libraries. P.S. and S.J. analyzed scRNA-seq data. S.J. additionally analyzed bulk transcriptomic data. A.D. was involved in data interpretation. M.L. and M.K.A. wrote the manuscript. M.K.A. additionally conceptualized and supervised the study.

## DECLARATION OF INTERESTS

The authors have declared that no conflict of interest exists.

## STAR METHODS

### Resource Availability

#### Lead contact

Further information and requests for resources and reagents should be directed to and will be fulfilled by the lead contact, Manish K. Aghi

#### Materials availability.

This study did not generate new unique reagents.

#### Data and code availability

- Deidentified human patient single-cell RNA-seq data reported in this study have been deposited at GEO and accession numbers are listed in the key resources table. They are publicly available as of the date of publication. In addition, summary statistics describing these data have been deposited at GEO and are publicly available as of the date of publication.
- All original code has been uploaded to FigShare and is publicly available as of the date of publication. Accession numbers are listed in the key resources table.
- Any additional information required to reanalyze the data reported in this paper is available from the lead contact upon request.

### Experimental Model and Subject Details

#### Cell lines.

Human GBM6 (Mayo Clinic), GBM43 (Mayo Clinic), DBTRG-05MG (ATCC), and U-251 (ATCC) cell lines were cultured at 37°C and 5% CO_2_ in Dulbecco’s modified Eagle’s medium (DMEM, Gibco) supplemented with 10% fetal bovine serum (FBS, Avantor) and 1% penicillin and streptomycin (P/S), GlutaMAX, non-essential amino acids (NEAA), and sodium pyruvate (Gibco). Murine GL261 (US National Cancer Institute) and BGL1 cell lines (gifted by Dr. Hideho Okada, Univ. Calif. San Francisco [UCSF]), as well as U-251 cells when used to generate U251cm, were cultured in Roswell Park Memorial Institute Medium 1640 (RPMI 1640) in similar conditions and using identical supplements, with the addition of 1% HEPES (Gibco). To generate GSC- containing neurospheres, GBM cells were grown in neurosphere media, consisting of DMEM/F12 (Gibco) supplemented with 20 ng/mL EGF (Peprotech), 20 ng/mL bFGF (Peprotech), and 2% GEM21/neuroplex (GeminiBio). All cell lines were passaged less than 10 times, screened bimonthly for mycoplasma, and validated every 6 months by Short Tandem Repeat (STR) analysis at the University of California Cell Culture Facility.

#### Mice

6-8 week-old female C57BL/6J, NuJ, and BalbC/J mice were purchased from the Jackson Laboratory. UBC-GFP mice were kindly provided by Dr. Harold Chapman, Dr. Martin C Valdearcos, and Dr. Suneil Koliwad (UCSF). All mice were housed in specific pathogen-free (SPF) conditions and standard environmental parameters (12:12 light:dark cycle; 10-15 air changes/hour; 30-70% humidity; 68-79°F) at the UCSF Helen Diller Cancer Research Building Animal Facility. In vivo vertebrate experiments were conducted using 8-12 week-old mice randomized to experimental groups, in accordance with UCSF IACUC protocol AN105170-02.

#### Human samples.

Human tumor samples were obtained from newly diagnosed IDHwt glioblastoma patients at time of initial surgical resection, following all guidelines stipulated by the UCSF Brain Tumor Research Center (BTRC) and in IRB #11-06160. Informed written consent was obtained from all patients preoperatively. Patient details, including age, sex, and experimental relevance, are provided in **Supplemental Table 1**.

### Method Details

#### Murine and human tissue harvest and dissociation

##### Tumor

Patient tumors were collected in serum-free Hibernate-A medium (Gibco) and transferred on ice to the laboratory for further processing. To harvest murine tumors, mice were first euthanized via CO2 administration (2L/min) in a closed chamber, followed by systemic perfusion with 10 mL of Dulbecco’s phosphate-buffered saline (DPBS; Gibco) through the left ventricle; the brain was subsequently isolated and the tumor resected using sterile surgical blades (Exel). To dissociate both human and murine tissue into single cell suspensions, tumors were washed twice with DPBS, cut into 1mm fragments, and incubated for 45 minutes at 37°C in 600 µL of dissociation buffer (0.74 U/µL collagenase IV [Gibco] + 5.3 U/µL deoxyribonuclease I [Worthington] in DPBS) per 100 mg tissue. Homogenized samples were then filtered through a 70 µm filter (Falcon) and resuspended in 1 mL ACK buffer (Gibco) for 2 minutes at room temperature (RT) to lyse erythrocytes. Final single cell suspensions were kept in DPBS on ice until downstream use.

##### Blood PBMCs

Patient and healthy control blood samples were collected in either EDTA- or heparin-coated tubes (BD) for subsequent neutrophil or T cell isolation, respectively. Murine blood was collected in EDTA-coated capillary tubes (RAM Scientific), either through saphenous vein draw in live mice or intracardiac draw in euthanized mice. Samples intended for flow cytometric analysis were additionally incubated in ACK buffer at a 1:10 ratio for 10 minutes at RT with intermittent agitation. Processed samples were kept on ice, with the exception of blood intended for neutrophil isolation, which was stored at RT to minimize thermal induction of neutrophil activation.

##### Systemic and skull bone marrow

To harvest systemic marrow, femurs were isolated from euthanized mice, their epiphyses were cut using surgical scissors (Excelta), and the internal marrow was flushed with DPBS using a 5 mL syringe (BD) fitted with a 25G needle (BD). Marrow contents were then passed through a 70 µm filter using a syringe plunger, and subsequently resuspended in 1 mL ACK buffer for 2 minutes at RT. To harvest skull bone marrow, calvaria were first isolated from euthanized mice by cutting around the occipital, parietal, and frontal bones using surgical scissors. The underlying dura was carefully removed using fine-toothed forceps (Integra). Skulls were then cut into 0.5-1.0 mm fragments, resuspended in 1 mL of dissociation buffer (as described earlier), and incubated for 45 minutes at 37°C with intermittent vortexing. The resulting mixture was passed through a 70 µm filter and resuspended in 1 mL ACK buffer for 2 minutes at RT. Final skull and systemic marrow samples were kept in DPBS on ice until use.

#### Staining for flow cytometric immunophenotyping and functional characterization

##### Staining protocol

All cell suspensions were processed and acquired immediately after harvesting, and antibodies were purchased from either BD or Biolegend. Samples were first stained with ZombieAqua or ZombieNIR viability dye according to manufacturer instructions. For apopotosis-related assays, Apotracker Green was also applied at this step at a working concentration of 0.16 µM. After washing with FACS buffer (2% FBS in DPBS), samples were then blocked for 10 minutes at 4°C in 10 µg/ml of either rat anti-mouse CD16/32 (clone 2.4G2; for murine samples) or Human Fc Block (Fc1; for human samples). In some cases, murine samples were alternatively blocked with anti-CD16/32-BV421 (190909) to enable characterization of this marker. Surface staining was otherwise completed following manufacturer recommendations. Where necessary, intracellular staining was performed using an eBioscience Foxp3/Transcription Factor Staining Buffer Set. Samples were acquired on an Attune NxT (ThermoFisher) flow cytometer, and data was subsequently analyzed with FlowJo software (Tree Star). All analyses were initiated with exclusion of cell doublets and dead cells.

##### Immune profiling

To characterize immune populations in human samples, cells were stained with fluorophore-conjugated anti-human antibodies against: CD3 (UCHT1), CD4 (OKT4), CD8 (SK1), CD11b (M1/70), CD11c (3.9), CD14 (M5E2), CD15 (W6D3), CD45 (2D1), CD56 (QA17A16), CD66b (6/40c), CD206 (15-2), MHCII/HLA-DR,P,Q (Tü39). For murine studies, fluorophore-conjugated anti-mouse antibodies were used: CD45 (I3/2.3, 30-F11), CD11b (M1/70), CD11c (N418), B220 (RA3-6B2), CD90.2 (30-H12), NK1.1 (S17016D), CD115/CSF1R (AFS98), F4/80 (BM8), CD117/c-Kit (2B8), Ly6G (1A8), CXCR2 (V48-2310, SA044G4), Ly6C (HK1.4), SiglecF (S17007L), Gr1 (RB6-8C5), CD3 (17A2), CD4 (GK1.5), CD8a (53-6.7), MHCII/HLA-IA,E (M5/114.15.2), CD206/MMR (C068C2), P2RY12 (S16007D), and CX3CR1 (SA011F11). CD45^+^ mouse hematopoietic cells were profiled into cKit^hi^ HSPCs, CD3^+^ T cells, CD11b^+^ myeloid cells, CD11b^-^B220^+^CD11c^+^MHCII^+^ pDCs, and CD11b^-^B220^-^CD11c^+^MHCII^+^ cDC1s. T cells were subdivided into CD8^-^CD4^+^ Th cells, CD8^+^CD4^-^ CTLs, CD8^+^CD4^+^ DNTs, and CD8^+^CD4^+^ double positive T cells. Myeloid cells were subdivided into cKit^hi^ precursors, Gr1^+^SiglecF^+^ eosinophils, CD11c^+^MHCII^+^CSF1R^lo^ cDC2s, P2Y12R^+^CX3CR1^+^ microglia, Gr1^+^Ly6G^lo/hi^CSF1R^-^ neutrophils, and CSF1R^+^/F480^+^ macrophages. Macrophages were further characterized as M1 (CD206^hi^MHCII^lo^), M2 (CD206^lo^MHCII^hi^), and/or inflammatory (Ly6C^+^), and neutrophils were defined as hybrid (CD11c^+^MHCII^+^), immature (CXCR2^-^Ly6G^lo/hi^), and/or mature (CXCR2^+^Ly6G^hi^).

##### T cell profiling

To evaluate activation in control-and patient-derived T cells cocultured with patient neutrophils, cells were stained with fluorophore-conjugated anti-human antibodies against: CD4 (SK3), CD8 (SK1), CD25 (M-A251), CD69 (FN50). For profiling of murine intratumoral T cell subsets, fluorophore-conjugated anti-mouse antibodies were used: CD45 (30-F11), CD3 (17A2), CD4 (GK1.5), CD8a (53-6.7), CD44 (IM7), CCR7 (4B12), GranzymeB (QA18A28; intracellular), CD62L (MEL-14), and CD69 (H1.2F3). CD3^+^CD4^+^CD8^-^ Th cells and CD3^+^CD4^-^CD8^+^ CTLs were individually identified as naïve CD62L^hi^CD44^-^ Tn’s, effector memory CD62L^lo^CD44^+^ Tem’s, and central memory CD62L^hi^CD44^+^ Tcm’s. Additionally, CTL activation was measured by CD69 and GranzymeB expression.

##### Neutrophil maturity and phenotype profiling

To identify immature and hybrid neutrophils among patient TANs and PBNs, human tumor and PBMC specimens were stained with fluorophore-conjugated anti-human antibodies against CD16 (3G8), CD66b (6/40c), CD15 (W6D3), MHCII/HLA-DR,P,Q (Tü39), CD49d (9F10), CD101 (BB27), CD10 (HI10a), CD3 (UCHT1), CD19 (HIB19), CD56 (HCD56), CD14 (M5E2), Siglec8 (7C9). For maturity analysis, after the exclusion of lineage-positive cells (CD3, CD19, CD56, CD14, Siglec8), CD66b^+^[CD16^+^/CD15^+^] neutrophils were assessed for their expression of MHCII and their internal composition: CD49d^hi^CD101^lo^ preNeus, CD49d^lo^CD101^hi^CD10^lo^ immNeus, and CD49d^lo^CD101^hi^CD10^hi^ matNeus. In murine studies – which included phenotypic induction experiments, migration experiments, and TAN/PBN profiling – cells were stained with fluorophore-conjugated anti-mouse antibodies against CD45 (I3/2.3, 30- F11), CD11b (M1/70), B220 (RA3-6B2), CD90.2 (30-H12), SiglecF (S7007L), NK1.1 (S17016D), CD117/c-Kit (2B8), CD16/32 (190909), CD11c (N418), MHCII/HLA-IA,E (M5/114.15.2), Gr1 (RB6-8C5), CD115/CSF1R (AFS98), Ly6G (1A8), F4/80 (BM8), and CXCR2 (V48-2310, SA044G4). Following exclusion of lineage-positive cells (SiglecF, NK1.1, CD90.2, B220), cKit^hi^CD16/32^+^ GMPs, CD11b^+^[CSF1R^+^/F480^+^] macrophages, and CD11b^+^Gr1^+^CSF1R^-^ neutrophils were identified. The latter were further stratified as immature CXCR2^-^Ly6G^lo/hi^ or mature CXCR2^+^Ly6G^hi^.

##### Glioma stemness characterization

Murine tumor stemness was characterized by staining mouse tumor samples with CD133 (315-2C11) and SOX2 (14A6A34). Glioma models were validated to express both markers *in vitro* prior to intracranial implantation.

#### Magnetic and fluorescence-activated cell sorting of human immune cells

For use in coculture studies, patient and control CD3^+^ T cells were isolated from PBMCs via negative magnetic selection (StemCell), following the manufacturer’s protocol. Similarly, PBNs were initially enriched via negative magnetic selection (StemCell). Both PBNs and TANs were subsequently purified from their respective samples (either enriched PBNs or tumor cell suspensions) via FACS. Cells were stained with fluorophore-conjugated anti-human CD45 (HI30), CD16 (3G8), and CD66b (6/40c), and live CD45^+^CD66b^+^CD16^hi^ neutrophils were sorted on a 4-laser 6-channel Sony SH800 cell sorter using ‘normal purity’ settings. For specimens intended for scRNA-seq analysis, sorted cells were collected in 5 mL round-bottom polystyrene tubes (Corning Falcon) prefilled with 10 µL of Protector RNase Inhibitor (Roche).

#### Fluorescence-activated cell sorting of mouse immune cells

To harvest murine BMNs for MHCII induction experiments, bone marrow isolates were stained with fluorophore-conjugated anti-mouse CD117/c-Kit (2B8), CD90.2 (30-H12), B220 (RA3-6B2), NK1.1 (S17016D), SiglecF (S17007L), CD115/CSF1R (AFS98), CD11b (M1/70), Gr1 (RB6-8C5), and CXCR2 (V48-2310). Following exclusion of HSPCs (cKit^hi^) and lineage-positive (CD90.2, B220, NK1.1, SiglecF, CSF1R) cells, immature CD11b^+^Gr1^+^CXCR2^-^ and mature CD11b^+^Gr1^+^CXCR2^+^ BMNs were sorted on a 4-laser 6-channel Sony SH800 cell sorter using ‘normal purity’ settings.

#### Imaging and analysis

##### Frozen section immunohistochemistry for TAN proximity to vasculature

Tissue obtained from the operating room was promptly suspended in 4% paraformaldehyde in DPBS for 12 hours, then transferred to a 30% sucrose w/v solution for 48 hours. Next, samples were submerged in Tissue-Plus Optimal Cooling Temperature (OCT) Compound TM (Fisher Scientific) and frozen at −80°C for >24 hours. OCT tissue blocks were sectioned into 10 µm slices using a Leica HM550 Cryostat and immediately adhered to Superfrost+ microscope slides (ThermoFisher), which were then refrozen at −80°C until staining.

Slides were fixed and stained in the typical fashion. Briefly, samples were rinsed with DPBS followed by blocking at RT for 1 hour in 5% goat serum (Jackson Immuno), 2% bovine serum albumin (Sigma Aldrich), and 0.3% Triton X-100 (Sigma Aldrich) in DPBS. Slides were incubated overnight at 4°C in anti-human primary polyclonal antibodies diluted in blocking buffer per manufacturer recommendations: anti-CD31 and either anti-MPO or anti-Nestin (Abcam). Samples were then rinsed with DPBS and incubated in secondary antibodies diluted in blocking buffer per manufacturer recommendations (Invitrogen) for 2h. The solution was aspirated and slides were allowed to air dry before mounting with DAPI (Southern Biotech) and coverslip (Fisher Scientific) application. Images were acquired on a Zeiss M1 fluorescent confocal microscope, and processed on ImageJ (Fiji).

##### Form factor analysis

Isolated TANs and PBNs were resuspended in DAPI and mounted onto slides prior to image acquisition on a Nikon Eclipse Ti-E epifluorescence microscope and subsequent analysis using ImageJ. As previously described, form factor or dendriticity was calculated as P^2^/(4*π*A), where P is cellular perimeter and A is area (Heffron and Mandell, 2005; Levi-Schaffer et al., 2000).

##### Cytocentrifugation and Wright-Giemsa histological staining

Isolated patient TANs/PBNs and murine BMNs were resuspended in supplemented RPMI and 25,000 cells were seeded per well into 8-well Nunc Lab-Tek chamber slides (Thermo Fisher). Slides were placed into a custom 3D-printed adapter (Castroagudin et al., 2016) and spun at 130 rcf for 5 minutes in a standard Eppendorf 5415 centrifuge. Wells were removed and samples were allowed to air-dry completely. Slides were then flooded with 1-2 mL of Wright-Giemsa stain (RICCA), followed 1 minute later by an equal volume of deionized water. After 3 minutes, slides were rinsed with deionized water and allowed to air dry before mounting with Cytoseal 60 (Epredia). Brightfield images were acquired on a Keyence BZ-X800 microscope using a Plan Apo 20x NA 0.75 WD 1mm objective. For nuclear segmentation analysis, nuclear lobulation was manually recorded and measurements were validated by a second independent observer.

#### Generation of neutrophil-and tumor-conditioned medias

##### Neutrophil-conditioned media

TANs and PBNs were cultured in non-treated 12-well plates (Genesee Scientific) for 12h in supplemented RPMI at a concentration of 10,000 cells/mL. Cells were then centrifuged and conditioned media collected. For purification, media was passed through a 0.45 µm Steriflip vacuum filtration unit (MilliporeSigma).

##### Tumor-conditioned media (TCM)

For generation of conditioned media from patient tumor specimens, viable CD45^-^ tumor cells were first enriched via FACS. Cells were then cultured for 12h in 10 mL of supplemented RPMI in non-treated 75 cm^2^ flasks (CellStar), at a concentration of 1.0E6 cells/mL. To generate condition media from immortalized tumor lines, 3.75E6 BGL1 and U251-related cells suspended in 28.125 mL of supplemented RPMI were seeded on to TC-treated 75 cm^2^ flasks (Greiner Bio-One CellStar) for 24h. Conversely, for GL261-conditioned media, 1.0E6 cells were first seeded in 10 mL of supplemented RPMI in TC-treated 75 cm^2^ flasks, then media was replaced after 72h with an additional 25 mL, and cells were allowed to incubate for 24 more hours. In all models, after their respective culture times, media was collected, centrifuged to remove cellular components, and further purified through 0.45 µm vacuum filtration.

#### Human neutrophil culture studies

##### Effects of tumor contact and TCM on neutrophil biology

To assess the effects of the tumor secretome on neutrophil viability, form factor, and MHCII expression, control and/or patient PBNs were cultured in TCM at a concentration of 150,000 cells/mL in non-treated 12-well plates; control samples were instead cultured in supplemented RPMI. In some experiments, PBNs were instead cultured directly with glioma cells (either U251 or autologous tumor); in these cases, 100,000 tumor cells were first seeded in 2 mL of supplemented RPMI, and media was replaced 12h later with a fresh 2 mL containing 300,000 PBNs. For blocking experiments, TCM was supplemented with 10 µg/mL anti-human anti-GM-CSF antibody (BVD2-23B6; Biolegend) or rat IgG2a κ (RTK2758; Biolegend) and incubated on ice for 30 minutes prior to use. Culture durations varied from 12h (form factor) to 96h with interval timepoints (viability, MHCII induction). For analysis, cells were first harvested via media collection and centrifugation, then imaged or assessed cytometrically, as previously described.

##### Effects of TANs and PBNs on tumor biology [NETosis]

We adapted a previously published murine NETosis assay to compare the effector capabilities of patient TANs and PBNs (McGill et al., 2021). Briefly, neutrophils were resuspended in supplemented RPMI with 0.05% v/v SYTOX Green (1:250 dilution in DMSO, Invitrogen) and either 50% v/v DMSO (Sigma Aldrich) or 50% v/v LPS (1:250 dilution in DMSO; Invitrogen eBioscience), at a concentration of 1.0E6 cells/mL. 200 µL of each suspension was plated per well of a non-treated 96-well flat-bottom plate (Costar) and incubated at 37°C/5% CO_2_ for 2h. Samples were harvested, washed and resuspended in FACS buffer, and acquired on an Attune NxT flow cytometer. NETosis induction per experimental condition was calculated as the ratio of SYTOX Green positivity in LPS-treated samples to that in vehicle (DMSO)-treated samples.

##### Effects of TANs and PBNs on tumor biology [glioma stem cell formation]

For neurosphere formation assays, GBM6, DBTRG_05MG, or GBM43 GSCs were seeded in 96-well non-treated plates with 200 µL of neurosphere media at a density of 24 cells/well, or in 12-well non-treated plates with 2 mL of neurosphere media at a density of 10,000 cells/well. For experimental conditions, cells were instead suspended in a 1:1 mixture of neurosphere media and either TAN- or PBN- conditioned media. In osteopontin-blocking assays, samples were treated with 5 ug/mL anti-human OPN (polyclonal; R&D) or normal goat IgG (polyclonal; R&D). Cultures were incubated for 24h, after which spheres were quantified per well.

##### Effects of TANs and PBNs on tumor biology [osteopontin ELISA]

Patient TAN- and PBN-conditioned media were assessed in technical triplicate for osteopontin production using an OPN ELISA (R&D Systems), on undiluted samples following manufacturer instructions.

##### Effects of TANs and PBNs on tumor biology [antigen processing capacity/DQ-OVA uptake]

To test whether neutrophils could proteolyze exogenous peptide antigen, TANs/PBNs were seeded into 96-well non-treated plates at a concentration of 50,000 cells/well in 200 µL of supplemented RPMI with 10 µg/mL of DQ-OVA (Invitrogen). Samples were incubated at 37°C/5% CO_2_ for 2 hours, then harvested, washed, and assessed by flow cytometry for green fluorescence.

##### Effects of TANs and PBNs on tumor biology [T cell interactions]

In experiments where neutrophils were initially activated, cells were resuspended at a concentration of 1.0E6 cells/mL in 1 µg/mL LPS in DPBS or pure DPBS, and incubated for 1 hour at 37°C/5% CO_2_ followed by 2 washes with DPBS. Next, cells were resuspended at the same concentration in supplemented RPMI with 10 µg/mL anti-human anti-MHCII/HLA-DR,P,Q (Tü39; Biolegend) or mouse IgG2a κ (MOPC-173; Biolegend), then incubated for 30 minutes at 37°C/5% CO_2_ before being diluted 6-fold in supplemental RPMI to a final concentration of 1.67E5 cells/mL. Meanwhile, control-or patient-derived T cells were first stained with CFSE (Biolegend) per the manufacturer’s protocol, then resuspended at a concentration of 8.33E5 cells/mL in supplemented RPMI with or without 10 uL/mL of Immunocult Human CD3/CD28 T Cell Activator (Stem Cell). Neutrophil and T cell suspensions were mixed in a 1:1 ratio and immediately plated in 96-well non-treated flat-bottom plates, at a density of 1.2E5 cells/well. Cultures were incubated for 72h at 37°C/5% CO_2_ with daily mixing by micropipetting and final analysis by flow cytometry.

#### Murine bone marrow neutrophil and mixed isolate culture studies

##### Effects of TCM on BMN maturation and viability

Mixed bone marrow isolates from C57BL/6J mice were cultured in 12-well non-treated plates at a seeding density of 500,000 cells/well in 2 mL of supplemented RPMI (controls) or GL261cm. Samples were incubated for 72h, and assessed for viability and relative preponderance of immature neutrophils by flow cytometry.

##### Effects of TCM on BMN hybrid polarization

To compare the phenotypic plasticity of different BMN subsets, sorted immature and mature murine neutrophils were cultured in 96-well non-treated flat-bottom plates at a seeding density of 50,000 cells/well in 200 µL of supplemented RPMI (controls) or syngeneic TCM (BGL1cm for Balb/cJ, GL261cm for C57BL/6J). Samples were incubated for 48h, after which they were assessed by histological staining, flow cytometry, and DQ-OVA uptake assays (as previously described).

##### Effects of TCM on BMN migration

Mixed murine bone marrow isolates from C57BL/6J mice were resuspended in supplemented RPMI and seeded onto 3.0 µm- pore PET-coated transparent 12-well transwell inserts (Falcon) at a concentration of 3.0E6 cells/well. Inserts were then placed into 12-well non-treated plates containing 1.5 mL of either prewarmed supplemented RPMI or GL261cm per well. Migration was allowed to continue for 16h at 37°C/5% CO_2_, after which cells from above and below the insert filter were collected and analyzed by flow cytometry.

##### Effects of TCM on skull neutrophil migration *ex vivo*

To assess the chemotactic effects of tumor-secreted proteins on skull neutrophils *ex vivo*, mouse calvaria were thoroughly rinsed with 3 washes of DPBS and cut sagitally into paired halves. Each half was placed, inferior side-down, into 2 mL of media in a 12-well non-treated plate; for each skull, one half was suspended in supplemented RPMI, and the other in syngeneic TCM (BGL1cm for Balb/cJ; GL261cm for C57BL/6J). Cells were allowed to egress for 48h at 37°C/5% CO_2_. At endpoint, cells were collected by harvesting media and thoroughly rinsing calvaria with DPBS, then assessed by flow cytometry for maturity status.

#### Bulk transcriptomic studies

##### Quantitative polymerase chain reactions for NANOG, OCT4

RNA was extracted from GSCs using an RNeasy Mini kit (Qiagen) and stored at −80°C until use. cDNA was created using qScript XLT cDNA Supermix (Quanta Bio), following the manufacturer’s protocol. cDNA was diluted to a constant concentration for all samples to ensure similar nucleic acid loading. Quantitative PCR was carried out using Power SYBR Green Master Mix (Applied Biosystems) on an Applied Biosystems StepOne Real-Time PCR cycler following manufacturer guidelines: 95°C for 10 min, followed by 40 cycles of 95°C/15 sec and 60°C/1 min. Ct values were calculated using the StepOne software (Applied Biosystems). Samples were prepared with three technical replicates per primer pair, using beta-actin as a control housekeeping gene.

##### Nanostring multiplex transcriptomic analysis

RNA was extracted from patient and murine tumor single cell suspensions using an RNeasy Mini kit (Qiagen) and stored at −80°C until use. A bioanalyzer was used to determine quantity and quality of the RNA sample. RNA (100ng) from each sample was hybridized with the Nanostring Myeloid v2 Profiling (human) and Mouse Tumor Signaling 360 (murine) codesets for 18 hours. 30 µl of the reaction was loaded into the nCounter cartridge and run on the nCounter SPRINT Profiler. Quality control and alignment of the raw data was completed using nSolver (Nanostring). Differential gene expression was carried out using the DeSeq2 package in R, followed by pathway expression analysis – as defined in the KEGG 2019 Human database – using the Enrichr pipeline. Raw files were simultaneously analyzed on the Rosalind online platform (OnRamp Bio) to calculate normalized gene expression counts, differentially expressed gene signifiance, and cell type scores. Geometric means for dendritic cell gene expression were manually computed; to identify target genes, the Nanostring probe annotation list was queried for all targets exclusively associated with dendritic cells and not other immune subsets. Visualizations (heatmaps, volcano plots) were generated on Microsoft Excel 365.

#### scRNA-seq of patient-matched TANs and PBNs

##### Sample loading

scRNA-seq was carried out using the 10x genomics platform.10- 15,000 patient TANs and PBNs were isolated by FACS into Protector RNase Inhibitor as described earlier, then validated for viability on a Countess automated cell counter. Cells were loaded on a Chromium Controller (10X Genomics) according to manufacturer instructions. The emulsion underwent reverse transcription immediately. cDNA isolation and library preparation were performed per manufacturer recommendations for processing granulocytes, including 2 extra cycles of cDNA amplification; library quality control was performed using Qubit and BioAnalyzer to determine the concentration and average size [520-570bp]. The libraries were sequenced on a NovaSeq platform at the UCSF sequencing core.

##### Quality Control and Pre-Processing

Feature-barcode matrices were generated using the 10X CellRanger pipeline, with “- -force-cells" used to include low-UMI barcodes. Feature-barcodes and count matrices subsequently underwent pre-processing and quality control in R (version 4.2.0) using Seurat (version 4.3.0). Cells with greater than 20% mitochondrial gene expression or expression of fewer than 200 genes were excluded. The LogNormalize, SelectVariableFeatures, and ScaleData functions were used to perform log normalization, variable feature selection, and scaling prior to dimensionality reduction using PCA (RunPCA) and UMAP (RunUMAP). A nearest neighbor graph was constructed and cells were clustered using the Louvain algorithm implemented through the FindNeighbors and FindClusters functions using a clustering resolution of 0.5. Other immune populations were removed from further analysis based on expression of CD3D/CD3E/CD4+ (T cells), CD79A/B+ (B cells), and CD14+ (monocytes). After pre-processing and quality control there were 7,463 cells (94.1%) x 16,849 genes in the blood sample and 11,831 cells (85.5%) x 19,913 genes in the tumor sample.

##### Integration

Integration of the samples from blood and tumor was performed by selecting variable features in each sample (SelectIntegrationFeatures), identifying anchors using CCA (FindIntegrationAnchors), and finally integrating the data (IntegrateData). Scaling, dimensionality reduction, and clustering was performed as outlined above after selecting “RNA” as the default assay for the integrated sample.

##### Module scores

Module scores were computed for genes sets for MHC class I/II presentation, cytotoxicity, and chemotaxis using the AddModuleScore function of Seurat, referencing gene sets described previously (Evrard et al., 2018).

##### Data visualizations

Heatmaps, UMAP plots, and feature plots were produced using the default visualization methods of Seurat. Volcano plots were produced using the EnhancedVolcano package (version 1.14.0) in R. Diffusion maps were produced using destiny (version 3.10.0) using the entire cells x genes matrix from the tumor sample as input.

##### Trajectory analysis (RNA velocity)

The BAM file outputs from the 10X CellRanger pipeline were converted to loom files using velocyto (version 0.17.17) and were then processed in scVelo (version 0.2.4) to generate estimates of RNA velocity using spliced and unspliced counts using the dynamical modeling setting with default parameters. Estimates of latent time and velocity arrows were computed and overlayed onto the UMAP embedding computed through the Seurat pre-processing pipeline.

##### Gene Ontology, GSEA, and KEGG pathway analysis

clusterProfiler (version 4.4.4) was used for all gene ontology (GO), GSEA, and KEGG pathway analysis. During gene ontology analysis, the top 50 genes by log2 fold change that were differentially expressed in each comparison group were utilized. The pathview package (version 1.36.1) was used to aid in visualizing expression of genes in KEGG pathways of interest.

##### Ligand-receptor analysis

CellChat (version 1.6.1) was used to analyze ligand-receptor interactions among clusters of the tumor neutrophil dataset. The whole default ligand-receptor database was utilized, and the average gene expression per cell group was calculated using the trimean method.

#### *Murine* in vivo *studies*

For all cranial surgical procedures, mice were anesthetized with 1.5% isoflurane (Dechra) and administered pre-and post-operative subcutaneous analgesics (0.05-0.1 mg/kg buprenorphine HCL [Buprenex]; 5-10 mg/kg meloxicam [Alloxate]). Body temperature was maintained intraoperatively by placing mice on a heating pad at 37°C. Prior to midline incision, mice were administered subcutaneous 0.25% bupivacaine HCL (Patterson Vetinerary), heads were shaved if necessary, and exposed scalps were sterilized with 70% EtOH and 7.5% povidone-iodine (Purdue Frederick). After surgery, incisions were closed with 7mm Reflex wound clips (CellPoint Scientific). Mice were monitored daily and euthanized at pre-determined endpoints or when at protocol humane endpoint (i.e., >15% weight loss or neurologic deficits).

##### Intracranial tumor implantation

150,000 luc^+^ BGL1 cells or 55,000 luc^+^ GL261 cells were resuspended in 2 µL of Hank’s buffered saline solution (HBSS; Gibco) and implanted intracranially into the right frontal lobes of Nu/BalbC or C57BL6 mice, respectively. Tumor inoculation was completed stereotactically (Stoelting) at 1.9 mm right lateral to bregma and at a depth of 3.0 mm below the dural surface, using a 10 µL micro-syringe fitted with a 26G 1” 12° PS4 needle (Hamilton) needle.

##### Bioluminescent imaging (BLI)

*In vivo* tumor growth was monitored through bioluminescence imaging on an IVIS Spectrum imaging system, with images taken every 3 days starting from the sixth day after tumorization; imaging was terminated per experimental group after the first mortality. Mice were first administered 100 µL of 30 mg/mL D-luciferin Potassium Salt in DPBS (GoldBio) intraperitoneally, then anesthetized with 2.5% isoflurane. Automatic exposure settings were used for image acquisition 10 minutes post-injection, and signal was analyzed using the Living Image 4.2 software (Caliper Life Sciences). Tumor size was measured as total flux (photons/s) in a region of interest (ROI) outlining the entire tumor, and in some analyses, was normalized per mouse to a starting BLI value.

##### αLy6G-mediated depletion of murine neutrophils

Neutrophil depletion was initiated 24h prior to tumor inoculation and sustained over the experimental duration through recurrent doses every 3 days. Mice were injected intraperitoneally with 200 µg of anti-mouse anti-Ly6G (1A8; BioXCell) or matched rat IgG2a κ isotype (2A3; BioXCell) at initiation, and 100 µg thereafter. Depletion efficacy was monitored systemically through weekly saphenous vein blood draws, and at endpoint intratumorally; specimens were analyzed by flow cytometry for neutrophil abundance.

##### Transplantation of CMFDA-stained and GFP^+^ skull flaps

Donor bone flaps were collected from harvested UBC-GFP and BalbC/J mouse skulls by cutting 3 and 4 mm-diameter circular cranial windows around the frontonasal and lambdoid sutures, respectively, at midline. BalbC/J flaps were then additionally each incubated in 48-well plates (Falcon) in 200 µL of 83.3 µg/mL CellTracker Green CMFDA dye (Invitrogen). To make the staining solution, 50 µg of lyophilized CMFDA was first reconstituted in 3 µL of DMSO, then diluted to a total volume of 600 µL with DPBS. Flaps were allowed to stain at RT with intermittent shaking for 10 minutes, then washed thoroughly 3x with DPBS to quench the reaction. Prepared UBC-GFP and stained BalbC/J bone flaps were stored in DPBS on ice, protected from light, until use.

Skull flap transplantation was then performed as previously described (Cugurra et al., 2021). Briefly, an electrical rotary micro-drill fitted with a 0.5 mm tungsten carbide bit (Fine Science Tools) was used to create 3 and 4 mm-diameter circular cranial windows around the frontonasal and lambdoid sutures, respectively, at midline. The skull was cooled intermittently with 0.9% sodium chloride solution (Henry Schein), and craniotomy was carefully performed with fine-toothed forceps so as to not damage the underlying meninges. Donor bone flaps were then placed within the surgical window, and manually fitted with surgical scissors as necessary. Finally, the transplant was fixed in place using cyanoacrylate glue (VWR) applied with a sterile cotton swab (Fisher Scientific).

##### Lethal irradiation of mouse skull bone marrow

To eliminate the egress of immune cells from murine skull bone marrow to the GBM TME, mouse heads were lethally irradiated 24h prior to tumor implantation. Awake mice were restrained in size-1 holding fixtures (Precision X-Ray) attached to lead head-exposure body shields (Precision X-Ray), then administered 13 Gy of split dose irradiation (6.5 + 6.5 Gy, 4 hour interval) on a Precision XRad 320 irradiator. Instrument settings were as follows: 50 cm platform distance to source, F2 filter set, 320.0 KV, 12.50 mA. Skull bone marrow was harvested at 2d, 7d, and 21d in non-tumor-treated mice to validate irradiation efficacy; the total number of viable CD45^+^ immune cells per skull was assessed by flow cytometry, and compared to non-irradiated controls.

##### Administration of AMD3100 to the skull

To induce myeloid cell egress from calvarial marrow, AMD3100 was administered as previously described on stereotactically restrained mice (Herisson et al., 2018). An electrical drill was used to thin the outer periosteum at 2 spots near bregma and 3 spots near lambda. 10 µL of 1 mg/mL AMD3100 (Abcam) in DPBS was drawn into a 10 µL Hamilton micro-syringe fitted with a 34G 0.375” blunt needle, and 2 µL of drug was injected into each of the predrilled spots, using the needle to poke the final hole and being careful to not penetrate the lower periosteum or dura. Each injection occurred over 60 seconds, and the needle was held in place for an additional 30 seconds to minimize reflux. After needle removal, bone wax (Ethicon) was immediately applied using a cotton swab.

### Quantification and Statistical Analysis

Statistical analyses were done using either Prism version 9.5.1 (GraphPad) or the Real Statistics Resource Pack software (Release 7.6; ©2013-2021, Charles Zaiontz) on Microsoft Excel 365. Comparisons between two groups were conducted using a non-parametric two-way Student’s t test; comparisons between multiple groups were conducted via one-way ANOVA with post-hoc Tukey contrasts for parametric variables, or Kruskal-Wallis with post-hoc pairwise Wilcox contrasts for non-parametric measures. Pearson correlation coefficients and adjusted R^2^ values were calculated for all linear regressions between continuous variables. Kaplan-Meier analysis was carried out for *in vivo* survival studies. Where appropriate, statistical outliers were removed if values met Grubbs’ criteria.

Study parameters and statistical measures are all explicated in the text, figures, and corresponding legends. These details include: number of replicates, exact P values, choice of mean or median for measure of center, and choice of SD or SEM to represent variance. Significance was defined as P<0.05, with Bonferroni correction applied for multiple comparisons.

Some figures were created with BioRender.com.

## SUPPLEMENTAL TABLE LEGENDS (EXCEL TABLES)

**Table S1 (related to Figure 1): Patient characteristics.** Newly diagnosed GBM patient samples collected from 2017-2022, with experimental applications listed.

**Table S2 (related to Figure 1): Nanostring Myeloid Signaling Panel, Patient TANs/PBNs.** Differential expression of myeloid gene expression in patient-matched TANs vs PBNs (n=3 pairs), with Nanostring annotations listed per gene.

**Table S3 (related to Figure 1): Nanostring Myeloid Signaling Panel, Volunteer PBNs.** Differential expression of myeloid gene expression in volunteer PBNs cultured in control (RPMI) or tumor-conditioned (U251cm) media (n=3/group).

**Table S4 (related to Figure 3): Nanostring Tumor Signaling Panel, CD8a^hi^ vs CD8a^lo^ BalbC/J Mice.** Differential expression of tumor signaling gene expression in the most-and least-CD8a enriched BGL1 GBM-bearing BalbC/J mice (n=3/group), which were treated with isotype and αLy6G, respectively.

**Table S5 (related to Figure 4): scRNA-seq of Patient Neutrophils, PBNs vs TANs.** All significantly (p<0.05) differentially expressed genes between patient-matched PBNs and TANs from the scRNA-seq dataset.

**Table S6 (related to Figure 4): scRNA-seq of Patient Neutrophils, TAN Subsets.** (*Sheet 1, “All TAN Clusters”*) Top 50 differentially expressed gene per TAN cluster from the scRNA-seq dataset, with fold-change values indicating expression relative to all other TANs. (*Sheet 2, “Cluster 3 vs 4”*) Significantly (p<0.05) differentially expressed genes between cluster 3 APC TANs and cluster 4 cytotoxic TANs. (*Sheet 3, “Cluster 4 vs PBNs”*) Significantly (p<0.05) differentially expressed genes between cluster 4 cytotoxic TANs and PBNs. If gene was also assayed in the Nanostring Myeloid Signaling panel comparing volunteer PBNs cultured in control (RPMI) vs tumor-conditioned (U251), a value of 1 is assigned in column G. For these concomitant genes, if the fold-change direction was identical and significant (p<0.05) in both assays, “YES” is assigned in column H.

**Table S7 (related to Figure 4): scRNA-seq of Patient Neutrophils, scVelo Dynamically Expressed Genes.** 264 top dynamically expressed genes across patient neutrophil populations, from an scVelo analysis of combined patient TANs and PBN scRNA-seq data. Clusters are organized from left to right corresponding to the RNA vectors shown in **Fig. 4I**, with the top 50 dynamic genes per cluster listed in descending expression order.

## Notes

### Competing Interest Statement

The authors have declared no competing interest.

