## Supplementary figures and images for "Glioblastoma induces the recruitment and differentiation of hybrid neutrophils from skull bone marrow"

### Supp. Fig. 1

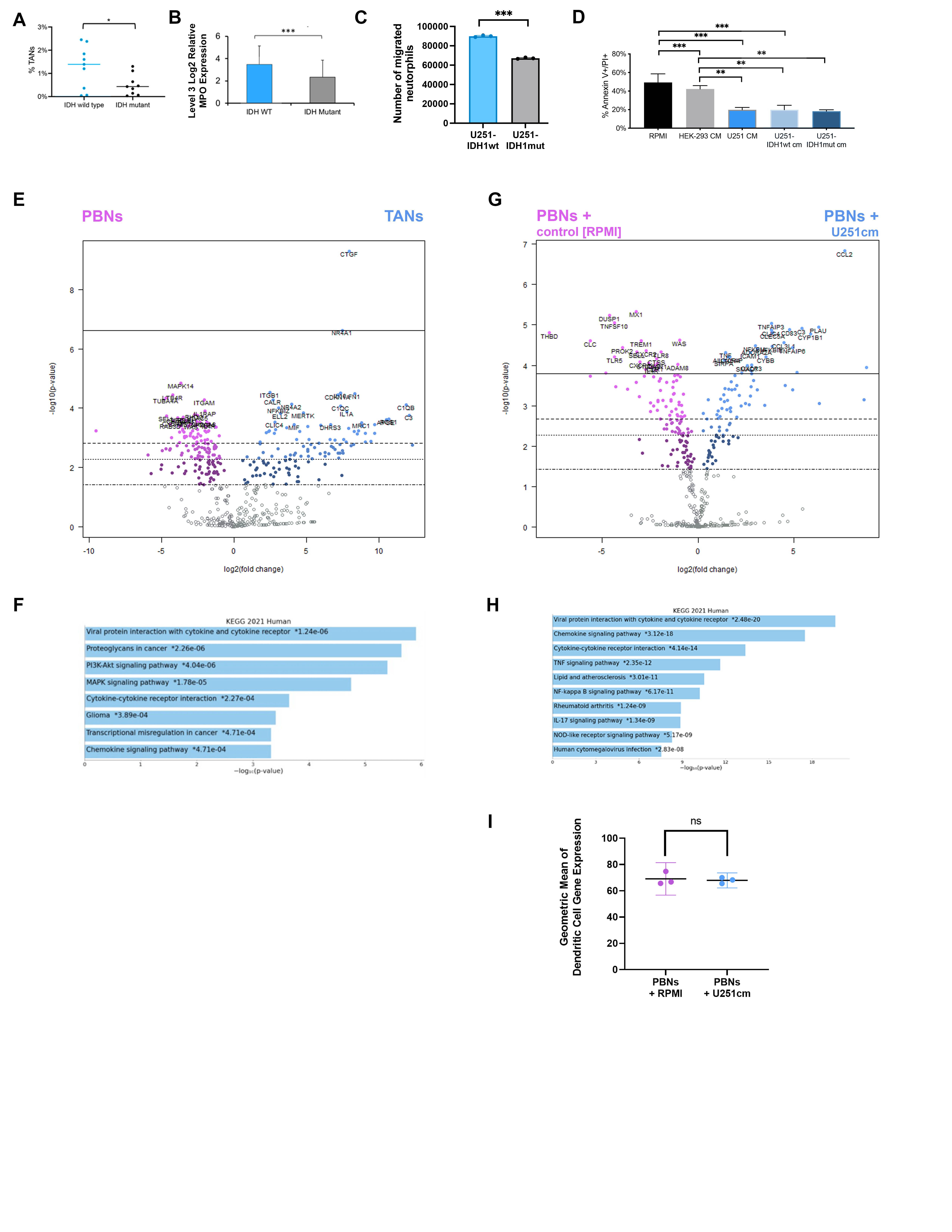

### Supp. Fig. 2

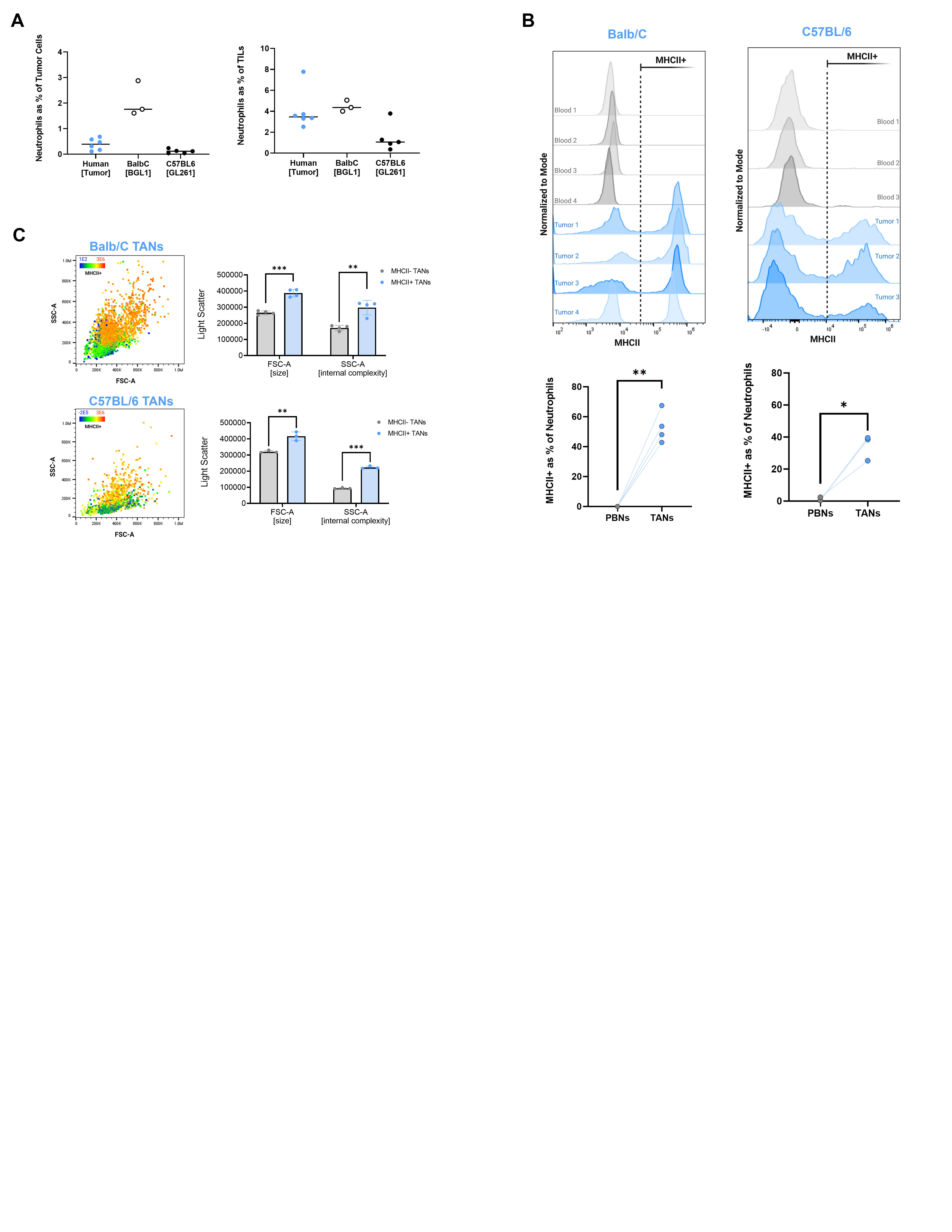

### Supp. Fig. 3

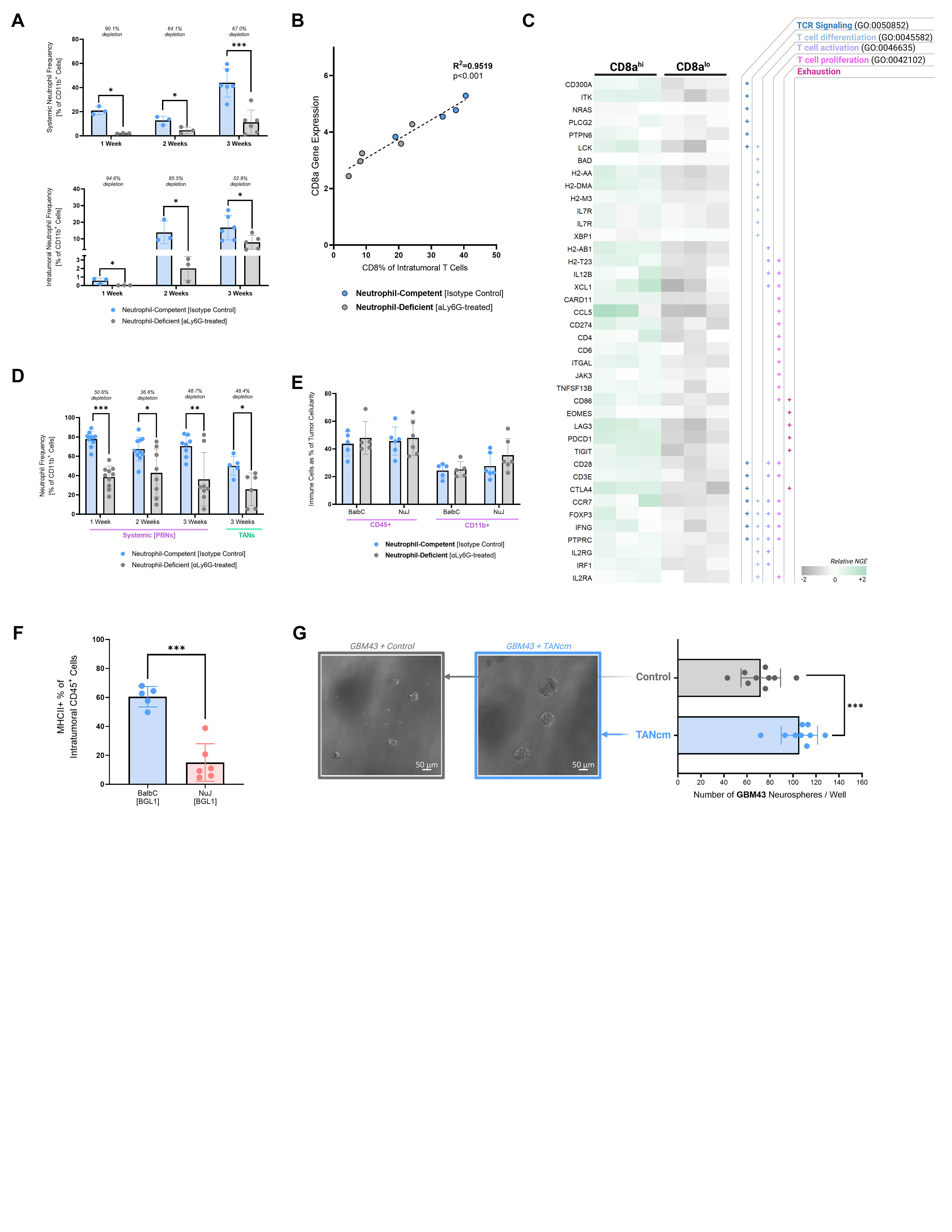

### Supp. Fig. 4

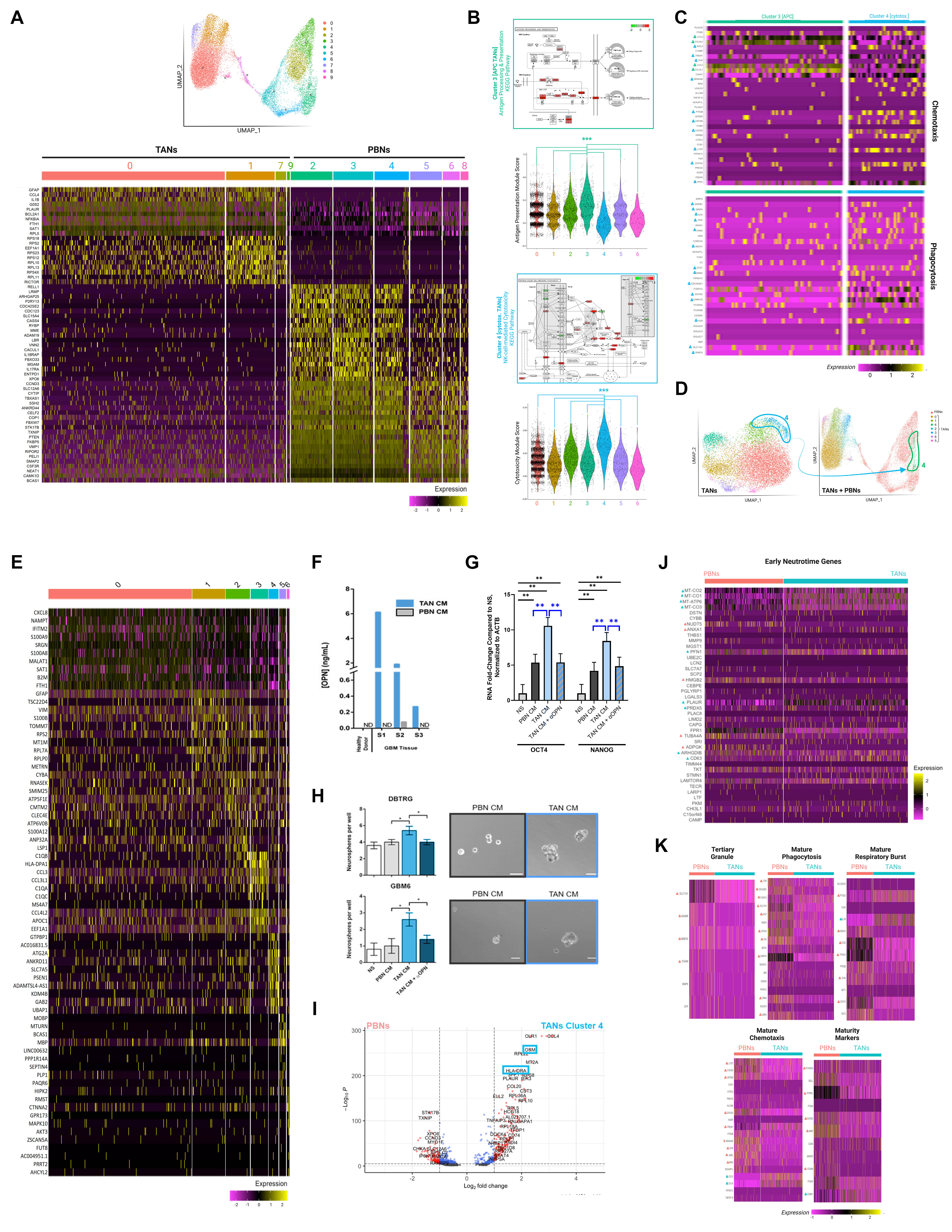

### Supp. Fig. 5

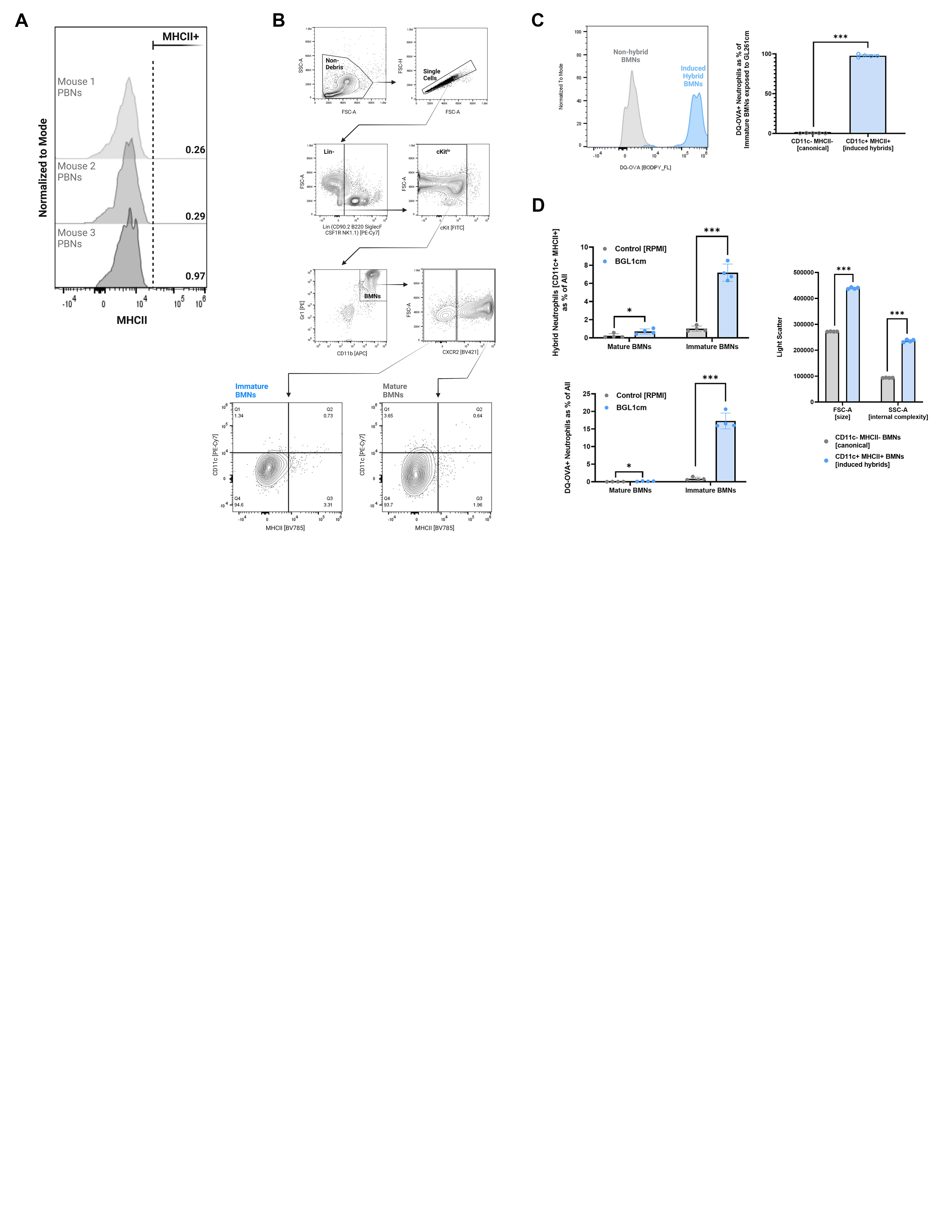

### Supp. Fig. 6

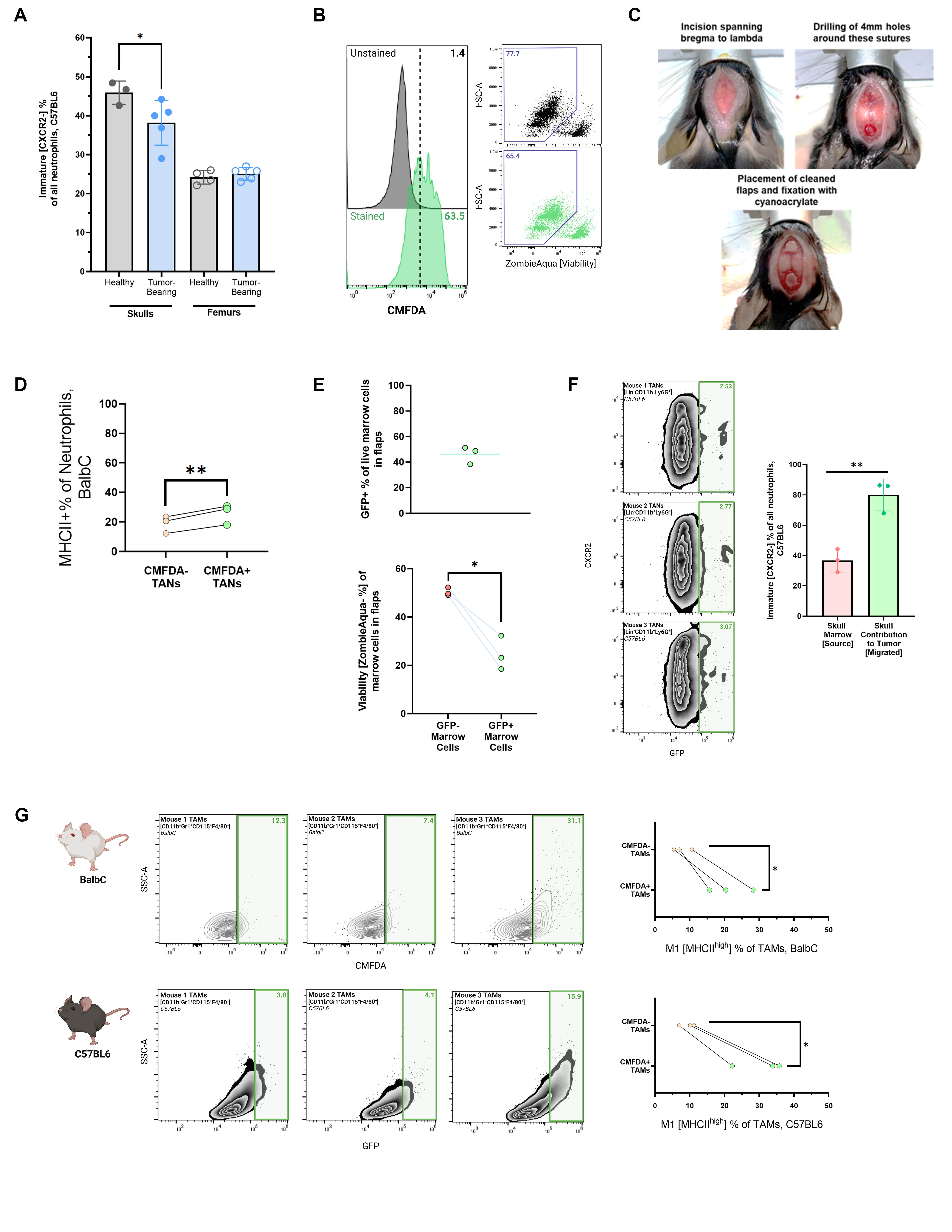

### Supp. Fig. 7

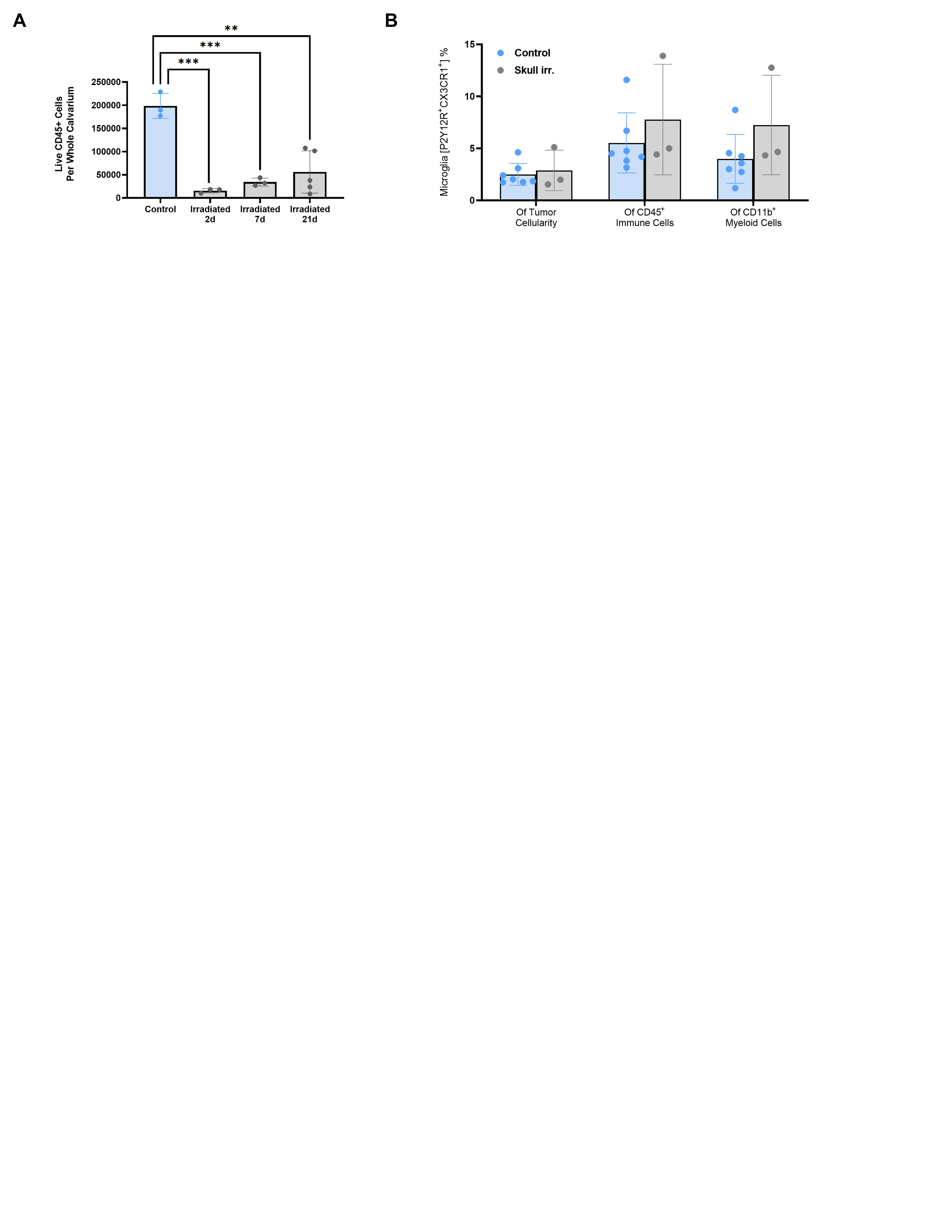
